# Biochemical deconstruction and reconstruction of Nuclear Matrix reveals the layers of nuclear organization

**DOI:** 10.1101/2023.01.28.525997

**Authors:** Ashish Bihani, Akshay K. Avvaru, Rakesh K. Mishra

## Abstract

Nuclear Matrix (NuMat) is the fraction of the eukaryotic nucleus insoluble to detergents and high-salt extractions that manifests as a pan-nuclear fiber-granule network. NuMat consists of ribonucleoprotein complexes, members of crucial nuclear functional modules, and DNA fragments. Although NuMat captures the organization of non-chromatin nuclear space, very little is known about component organization within NuMat. To understand the organization of NuMat components, we subfractionated it with increasing concentrations of the chaotrope Guanidinium Hydrochloride (GdnHCl) and analyzed the proteomic makeup of the fractions. We observe that the solubilization of proteins at different concentrations of GdnHCl is finite and independent of the broad biophysical properties of the protein sequences. Looking at the extraction pattern of the Nuclear Envelope and Nuclear Pore Complex, we surmise that this fractionation represents easily-solubilized/loosely-bound and difficultly-solubilized/tightly-bound components of NuMat. Microscopic analyses of the localization of key NuMat proteins across sequential GdnHCl extractions of *in situ* NuMat further elaborate on the divergent extraction patterns. Furthermore, we solubilized NuMat in 8M GdnHCl and upon removal of GdnHCl through dialysis, *en masse* renaturation leads to RNA-dependent self-assembly of fibrous structures. The major proteome component of the self-assembled fibers comes from the difficultly-solubilized, tightly-bound component. This fractionation of the NuMat reveals different organizational levels within it which may reflect the structural and functional organization of nuclear architecture.

## INTRODUCTION

The eukaryotic nucleus is a crowded milieu that accommodates, organizes, and regulates the genome – acting as the dynamic instructional repository of the cell (Misteli, 2005). To understand how the genome is managed, it is necessary to understand the architecture surrounding it. Biochemical investigations into the nuclear ultrastructure with different clearing buffers yielded different residual structures, allowing exploration of different aspects of the nuclear interior (reviewed in (Nelson et al., 1986)). Interestingly, extraction of nuclei with 0.4 M NaCl and 1.4 M NaCl leads to the solubilization of the chromatin. This reveals a granular-fibrous structure, stipulated at the time, to be made up of precursors of ribosomal subunits (Georgiev, 1967; Georgiev and Chentsov, 1962; Smetana et al., 1963). Treatment of such preparations with DNases solubilize more material and make the underlying network better visible, albeit highly compacted (Shankar Narayan et al., 1967). Through electron microscopy, this fiber network was shown to be the skeleton of the nucleus and was termed the Nuclear Matrix (Berezney and Coffey, 1974; Zbarsky, 1998; Zbarsky and Perevoshchikova, 1948) shortened as NuMat (Kallappagoudar et al., 2010).

NuMat is essentially the residual fraction after DNase I treatment and salt-detergent wash of nuclei, contains thousands of proteins (Engelke et al., 2014; Kallappagoudar et al., 2010; Sureka and Mishra, 2021), some of them complexing various kinds of nuclear RNAs (Nakagawa and Prasanth, 2011; Pathak et al., 2013; Zheng et al., 2010), and fragments of DNA that serve as attachment points for the chromatin to the matrix (Pathak et al., 2014). Using fluorescence microscopy and biochemical characterization, many crucial functions of the nucleus, such as transcription, replication, splicing, DNA repair, *etc.,* have been shown to use NuMat as a substrate (Berezney et al., 1995; Cook et al., 1999; Hozák et al., 1993; Koehler and Hanawalt, 1996; Zeitlin et al., 1987). A huge fraction of these components is also retained in an analogous structure derived from Mitotic chromosomes, termed Mitotic Chromosomal Scaffold, and is transmitted to the daughter cells (Adolph et al., 1977; Sureka et al., 2018).

Despite the extensive biochemical characterization, proteomics, RNA/DNA sequencing, and microscopic studies of NuMat, we do not know the organization of these components within the NuMat fibers, the mechanism of their assembly, or if they represent true *in vivo* architecture in any way (Pederson, 2000). We consider three possibilities of NuMat organization: I) NuMat is composed of different kinds of fibers, e.g., Motor protein fibers (Actin-myosin, Tubulin-Kinesin-Dynein), Intermediate Filaments (Lamin, Spectrin), RNP fibers (nucleolar proteins, ribosomal subunits, hnRNPs) (He et al., 1990; Tan et al., 2000), supporting architectural proteins (Megator, Topoisomerase II, Cohesins, Condensins, NuMa), *etc*.; II) The nucleus is divided into membrane-less compartments without a uniform underlying network based solely on liquid-liquid phase separation relationships of different protein-RNA complexes. The proteins may stick to their neighbours due to a sudden change in the environment as we remove the chromatin and other buffering agents during NuMat preparation, and form fibrous aggregates that are visible to us in form of NuMat; III) The fibers are aggregates of interchromatin RNPs forming channels for the transport of RNAs – which would form a random aggregate (Razin and Gromova, 1995). Either way, the dissection of NuMat fibers contains spatial information, either the structural core and the sticky-functional fraction forming the interface with the chromatin, or an ordered aggregate reflecting the spatial arrangement of biomolecules in subnuclear compartments.

To dissect NuMat organization, we use a well-known chaotrope, Guanidinium Hydrochloride (GdnHCl), which interferes with both electrostatic and hydrophobic interactions of protein folds by forming solvation stacks (Lim et al., 2009; Raskar et al., 2019), and hence, causes proteins to denature and solubilize better than most denaturants. We reasoned that GdnHCl would readily dissolve aggregates and loosely bound proteins, leaving behind the structural elements. We have performed a sequential extraction with increasing concentrations of GdnHCl, where the gross morphology of nuclei remains unperturbed. A fraction of NuMat protein content is solubilized in a finite time for each concentration of GdnHCl. Through proteomics of different GdnHCl fractions, we observe that NuMat proteins are differentially extracted irrespective of their biophysical properties. We also observe that the difficult-to-solubilize proteins have a propensity for RNA-dependent reassembly. Based on these observations, we posit that NuMat is a hierarchical structure capable of ultrastructural self-assembly, containing information about subnuclear organization and function.

## RESULTS

NuMat captures the non-chromatin architecture of the eukaryotic nucleus. To further understand the organization of components within NuMat ultrastructure, differential solubilization and subfractionation is needed. However, any further subfractions of NuMat have not been characterized. The filamentous architecture of NuMat has been observed to be eliminated by proteases or Sodium Deoxycholate treatments (Agutter and Richardson, 1980), while treatment with RNAse causes the fibrous structures in the nuclear interior to collapse (Comings and Okada, 1976; He et al., 1990). RNAse treatment also disrupts the overall chromatin organization (Barutcu et al., 2019; Nickerson et al., 1989). On the other hand, various salts/detergents do not show any significant differential solubilization (Engelke et al., 2014). We need a gradual biochemical extraction method. Therefore, we attempted to extract our NuMat prep (Fig EV1) with two chaotropes, Urea and GdnHCl. While extraction with different concentrations of Urea shows no solubilization (Fig EV2 a), buffers with increasing concentrations of GdnHCl are able to partially solubilize NuMat (Fig EV2 b). We used 1M, 2M, 3M, and 4M concentrations of GdnHCl to test and optimize fractionation. We show that incubation with GdnHCl extraction buffer of a particular concentration is able to solubilize a distinct set of proteins at that concentration with no further extraction happening upon longer incubation (Fig EV3). This is a strong indication that NuMat fibers do not result from random clumping of non-chromatin proteins. The finite extraction at each concentration enabled us to extract NuMat preps within a sequential manner (experimental schematic in Fig 1 a, gel profile in Fig EV4). The sequential extraction of NuMat yielded 6 fractions, labelled with the respective GdnHCl concentration. As shown in Fig EV4, the sequential extraction fractionation shows interesting solubilization patterns. Some proteins are solubilized by low ionic strength GdnHCl buffers (1M, 2M) while others remain insoluble (P). Additionally, it appears that despite exhaustive extraction at each step, several proteins repeatedly appear across fractions.

**Figure 1:**
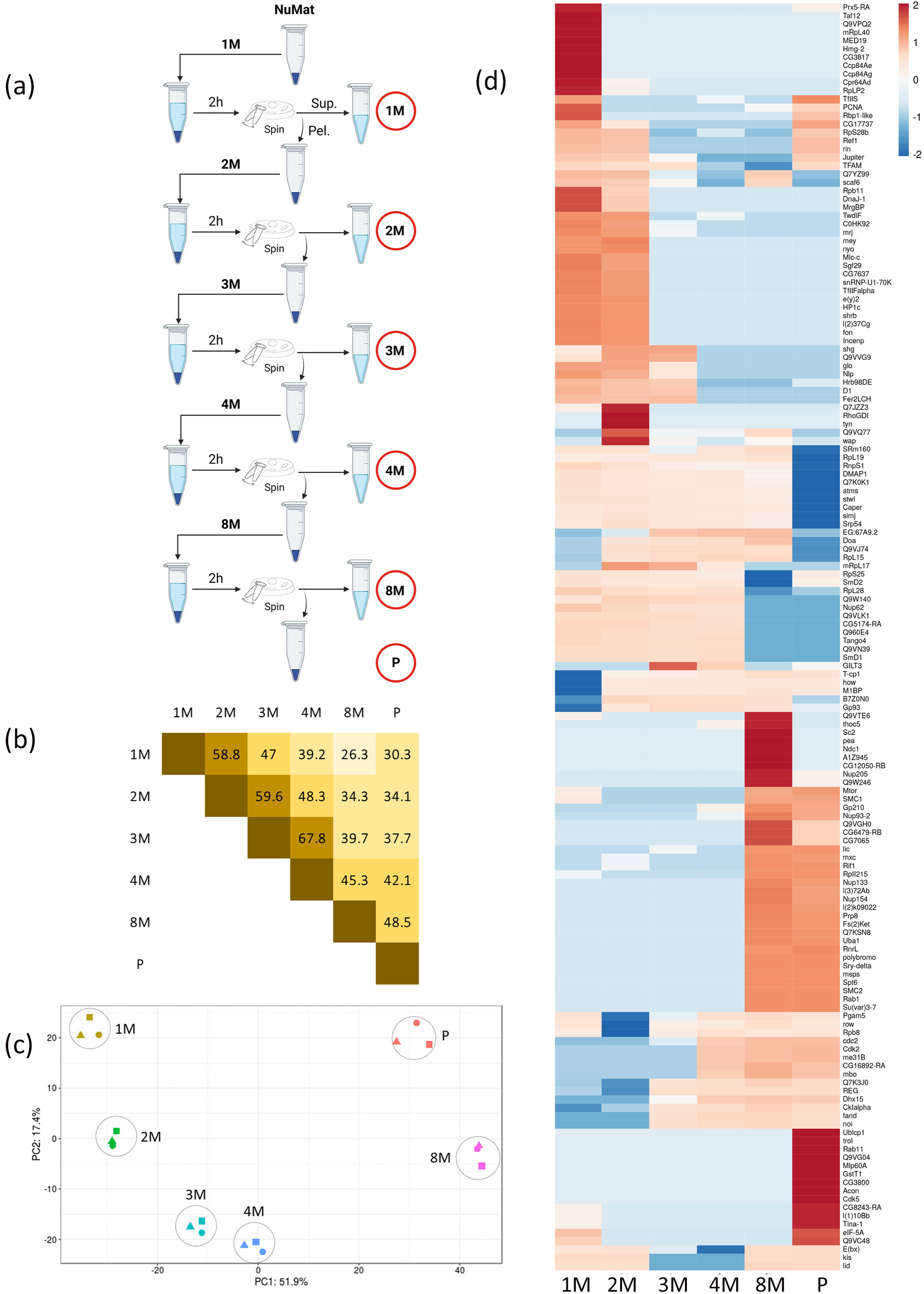
NuMat proteins show differential enrichment in different GdnHCl extraction fractions. **(a)** For sequential extraction of NuMat with different GdnHCl concentrations the procedure is as described in the schematic; NuMat is resuspended in extraction buffer of the mentioned GdnHCl concentration, incubated for 2h at 25° and then pelleted down. The supernatant is taken as extract while the pellet is resuspended in the extraction buffer containing a higher concentration of GdnHCl, repeated till 8M. The 1M, 2M, 3M, 4M, and 8M extracts and the final pellet (P) are subjected to proteomics (marked with red circles). **(b)** Pairwise percentage overlap for NuMat fractions. Shown here in a 6×6 matrix is the overlap of each fraction with another fraction, each box corresponding to the notation of two concentrations contains the percentage overlap of protein lists for those two fractions. The higher the extent of overlap, the more intense the color of the box. We see that the extraction profiles of fractions closer in terms of GdnHCl concentration overlap better. **(c)** Replicate correlation for all replicates across all fractions in the sequential extraction of *Drosophila* embryo NuMat and with different concentrations of GdnHCl. Each dot represents one sample (one replicate of one fraction, as labelled). The separation between dots denotes the difference between them while dots clustering together have a similar profile. Replicates of each fraction cluster together well and are distinct from other fractions **(d)** Quantitative enrichment profile of proteins across the fractions of NuMat. LFQ intensities of GdnHCl fractions of NuMat were compared using LFQ-analyst, filtered by fold change (>25log_2_-fold, p<0.001) without imputation and Benjamini-Hochberg FDR correction. Fraction-wise relative enrichment scores of selected proteins were calculated using DEP and plotted in a heatmap using ClustVis. Columns represent fractions (1M, 2M, 3M, 4M, 8M, and P) and rows represent individual proteins. The red color shows the enrichment of proteins and the blue color shows their depletion. We see a huge number of proteins are highly differentially enriched in different fractions and there is a stark division between higher-concentration and lower-concentration extractions.

### Proteomics of GdnHCl fractions of NuMat

To understand the GdnHCl solubility patterns of NuMat proteins, we performed proteomics on the GdnHCl extraction fractions of the NuMat (1M, 2M, 3M, 4M, 8M, P). Summing up all the proteins across different fractions, 908 proteins were identified (identification parameters in Supplementary Table 1, and identifications listed in Supplementary Table 2 sheet 1). Pairwise comparison of the protein lists of fractions shows that there is a significantly higher overlap between the proteins that are extracted at proximal concentrations (*e.g.*, 1M and 2M) and a smaller overlap at distant concentrations (*e.g.*, 1M and 8M), implying that the extraction is gradual (Fig 1 b, Supplementary Table 2 sheet 2). We consider the proteins solubilized with low ionic concentration GdnHCl buffers (1M-2M) as easily-solubilized and the proteins only solubilized with 8M GdnHCl or not solubilized even by 8M GdnHCl (8M-P) as difficultly solubilized.

The sets 1M-2M and 8M-P have large numbers of unique proteins while the set 3M-4M does not, *e.g.*, 1M-2M: 1M (55), 2M (46), 1M 2M (46), and 8M-P: 8M (153), P (57) 8M P (88) (Fig EV5, Supplementary Table 3). The Principle Component Analysis (PCA) of the quantitative proteome profiles of all replicates of all 6 fractions shows that the proteome intensity profile of the replicates of individual fractions are similar, and the fractions are distinct from each other. Also, a bigger divergence is seen between easily-solubilized and difficultly-solubilized proteins (Fig 1 c, PCA input data in Supplementary Table 4). We compared the quantitative profiles of the GdnHCl fractions by doing clustering analysis on differentially enriched proteins (p<0.001) with a high fold change (>25log_2_), red indicating a heavily enriched protein and blue indicating a heavily depleted protein (Fig 1 d). The heavily differentially enriched proteins make up ∼16.5% of the total proteins detected (150 out of 908) (Supplementary Table 5 sheet 1). Two major clusters of heavily enriched proteins emerge in this analysis, which represent classes of proteins that are extremely easy to solubilize (1M-2M) and proteins that are extremely difficult to solubilize (8M-P) (Fig 1 c). We reason that the differential solubility of these proteins is because of their broad biophysical affinity to a particular GdnHCl concentration. This could be due to biophysical properties intrinsic to their amino acid sequence or due to extrinsic factors like macromolecular interactions or post-translational modifications – dictating the manner of their spatial organization or packing in NuMat. However, we observe that the protein sets belonging to different extraction fractions do not show any significant enrichment in biophysical properties intrinsic to the amino acid sequences, *e.g.,* molecular weight, isoelectric point, intrinsic disorder, and percentage non-polar amino acids (Fig EV6, Supplementary Table 6).

Since the extractions are finite (Fig EV3), and yet the same proteins are extracted in multiple fractions, we surmise that there are multiple biophysical subpopulations of the same protein, leading to their differential extraction or solubilization. This becomes evident looking at ubiquitously-solubilized proteins. Comparing qualitative profiles of all fractions together, we observe that 132 proteins are common in all fractions (Fig EV5). In terms of LFQ too, a significant number of proteins (∼11.3%, 103 out of 908) are ubiquitous or display very low fold change (<2), which is statistically insignificant (p>0.05) (Supplementary Table 5 sheet 2). In addition, a large number of proteins show extraction enrichment in different extraction concentration ranges (say 1M and 8M). Such extraction patterns could be considered spurious if they were seen to happen with one or two outliers. However, a large number of proteins show bimodal or multimodal extractions, which implies that the individual proteins have multiple biophysical subpopulations due to modifications or interactions with other moieties.

### GdnHCl extraction of *in situ* NuMat preparation

NuMat prepared from purified *Drosophila* nuclei is very sticky and does not lend well to confocal microscopy. We prepared *in situ* NuMat from intact uncrosslinked fly embryos with a slightly modified protocol (Pathak et al., 2022a, 2022b). This method also uses salt extraction and DNase treatment to remove chromatin and chromatin-associated proteins from uncrosslinked nuclei, and hence replicates the biochemistry of NuMat in a microscopy-friendly manner (Pathak et al., 2022b). We extracted the *in situ* NuMats with GdnHCl extraction buffers in a sequential manner and found that the extracted embryo cage remains stable till 3M GdnHCl extraction and can be used to assess and further understand the GdnHCl extraction patterns of NuMat proteins. We looked at the extraction pattern of easy-to-solubilize proteins (Heterochromatin Protein 1 a (HP1a), Polycomb (Pc)), difficult-to-solubilize proteins (Megator/Tpr and RNA Pol II subunit 215 (RNAP2)), and ubiquitously-solubilized proteins (Lamin Dm0 (Lam), Fibrillarin (Fib), GAGA-associated factor (GAF)/Trl) in the uncrosslinked *in situ* NuMats obtained after each extraction step. Based on the distribution of these proteins in nuclear space, the extraction trends observed here are similar to those observed in extraction fraction proteomics (Fig 2). The spatial visualization of the extraction steps supplies additional information. Hp1 and Pc remain only in a few foci upon NuMat preparation and are extracted mostly in low GdnHCl (1M/2M) extractions (Fig 2 a, and b). GAF is extracted in a gradual manner (Fig 2 c). Megator and Lamin have primarily peripheral localization and they hold their place even after 3M GdnHCl extraction (Fig 2 d, and g, Fig EV7). The nucleolar foci of Fibrillarin remain strong through NuMat preparation and 1M GdnHCl extraction but their morphology starts to change as we go to higher concentrations (Fig 2 e). RNAP2, remains tightly bound (Fig 2 f). The LFQ intensity data for these particular proteins is shown in a line diagram (Fig EV8).

**Figure 2:**
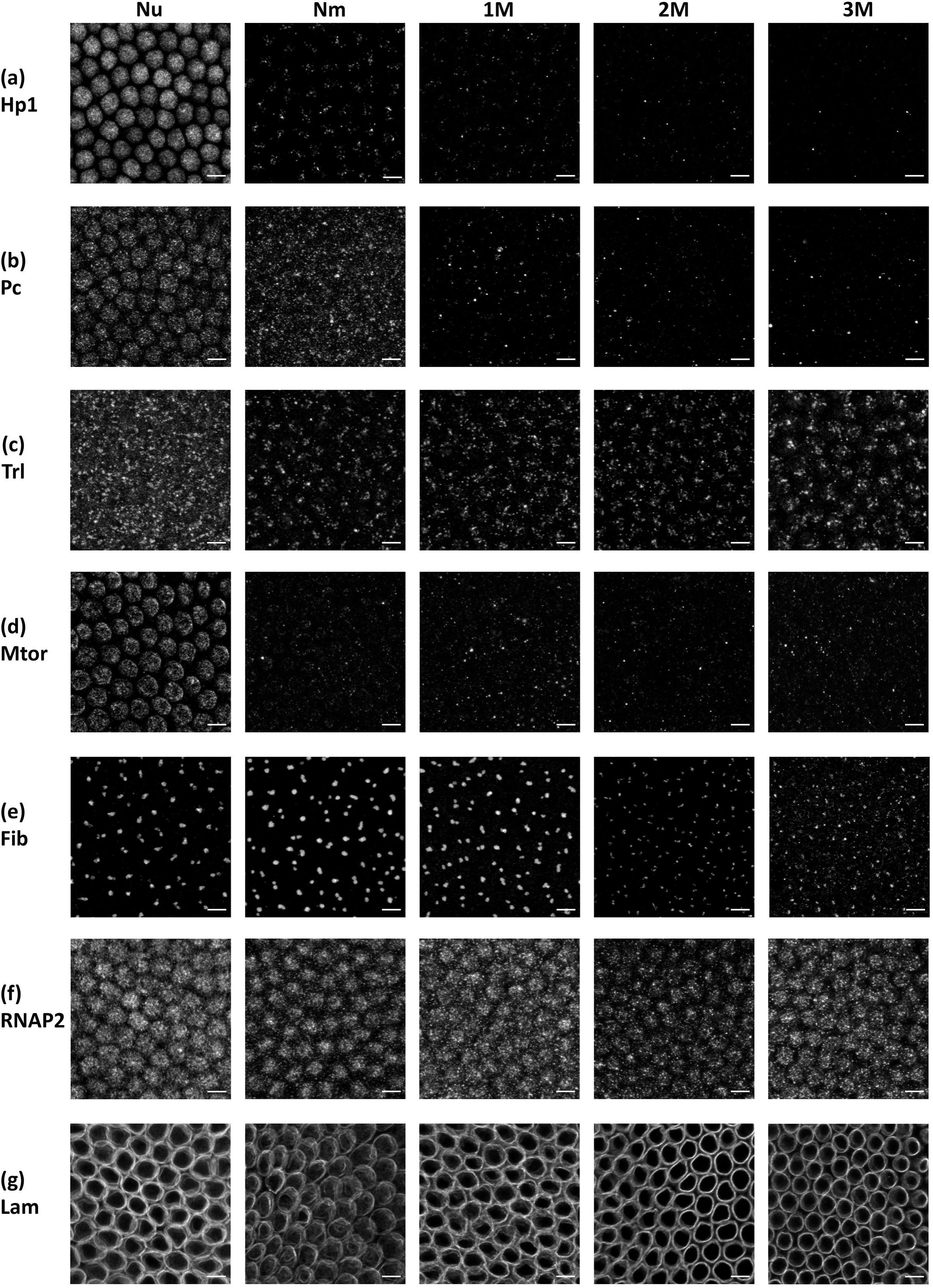
Extraction patterns of well-known NuMat proteins in *in situ* NuMat follow a trend similar to proteomics. Shown here are images of *Drosophila* embryos at the 13^th^-15^th^ stage (∼2h), at100x magnification and 3X zoom (**Nu** for Nuclei, **Nm** for NuMat, and **1M, 2M, 3M** extracts). Each row shows the sum projection of a sheet of nuclei immunostained for: **(a)** Hp1a **(b)** Polycomb **(c)** Trithorax-like **(d)** Megator **(e)** Fibrillarin **(f)** RNAP2 **(g)** Lamin; in greyscale to assess the extent of extraction for the sequentially extracted samples. Scale bar 5 µm in all images.

Fibrillarin and HP1 have very different behaviors as HP1 is extracted with the NuMat preparation from nuclei and in low-concentration GdnHCl washes while subpopulations of Fibrillarin get extracted at all GdnHCl concentrations. But both proteins are highly segregated scaffold proteins, building compact liquid-like environments in their respective compartments by multivalent interactions and association with nucleic acids (Keenen et al., 2021; Lafontaine, 2019). In general, it is worth looking at how these differentially extracted subpopulations of proteins form complex compartments, that get differentially extracted in such a starkly distinct manner (as seen in Fig 2 a, and e). While much is known about the roles of both GAF and RNAP2 in a regulatory and mechanistic sense, their dynamics in the subnuclear space needs further exploration. However, both proteins appear to constitute a dense distribution of foci across the nuclear interior (Fig 2 c, and f). The differential extraction data suggests that while GAF is a protein with many extraction modalities, RNAP2 remains unextracted. These observations elaborate that different proteins might be showing particular extraction patterns due to their biophysical status and their engagement with various subnuclear compartments. This also establishes *in situ* NuMat as a good platform for further biochemical and genetic investigations of solubility patterns of individual proteins.

### Renaturation of GdnHCl solubilized NuMat proteins

Homotypic or heterotypic self-association among nuclear proteins is crucial for compartment formation in nuclei. While recent developments in the study of biomolecular condensates provide valuable tools to look at self-association among soluble proteins, similar studies are difficult to do for insoluble endogenous proteins. To assess if NuMat proteins have any propensity for self-assembled ultrastructure formation, we dissolved the whole NuMat in 8M GdnHCl solution and cleared the undissolved fraction of debris (experimental schematic in Fig 3 a). The supernatant does not have any complex structure (Fig 3 b). Upon GdnHCl removal through dialysis, imaging the resultant sample with a transmission electron microscope shows the formation of networks of fibers (Fig 3 c). This shows that NuMat proteins can self-assemble into higher-order ultrastructures. The renatured fibers do not collapse when centrifuged at 21000xg (Fig EV9 a, and b). These fibers also form if GdnHCl concentration is reduced through dilution, also diluting the protein content drastically and therefore, are not dependent on protein concentrations (Fig EV9 c). Treatment of renatured NuMat with RNAse leads to the absence of fibers (Fig 3 d). We also prepared NuMat with RNAse treatment alongside DNase treatment. The renaturation of proteins from this prep also leads to many proteins becoming insoluble but they do not form fibers (Fig 3 e), implying that nuclear RNA is an important driver of the ultrastructure forming self-assembly of NuMat proteins.

**Figure 3:**
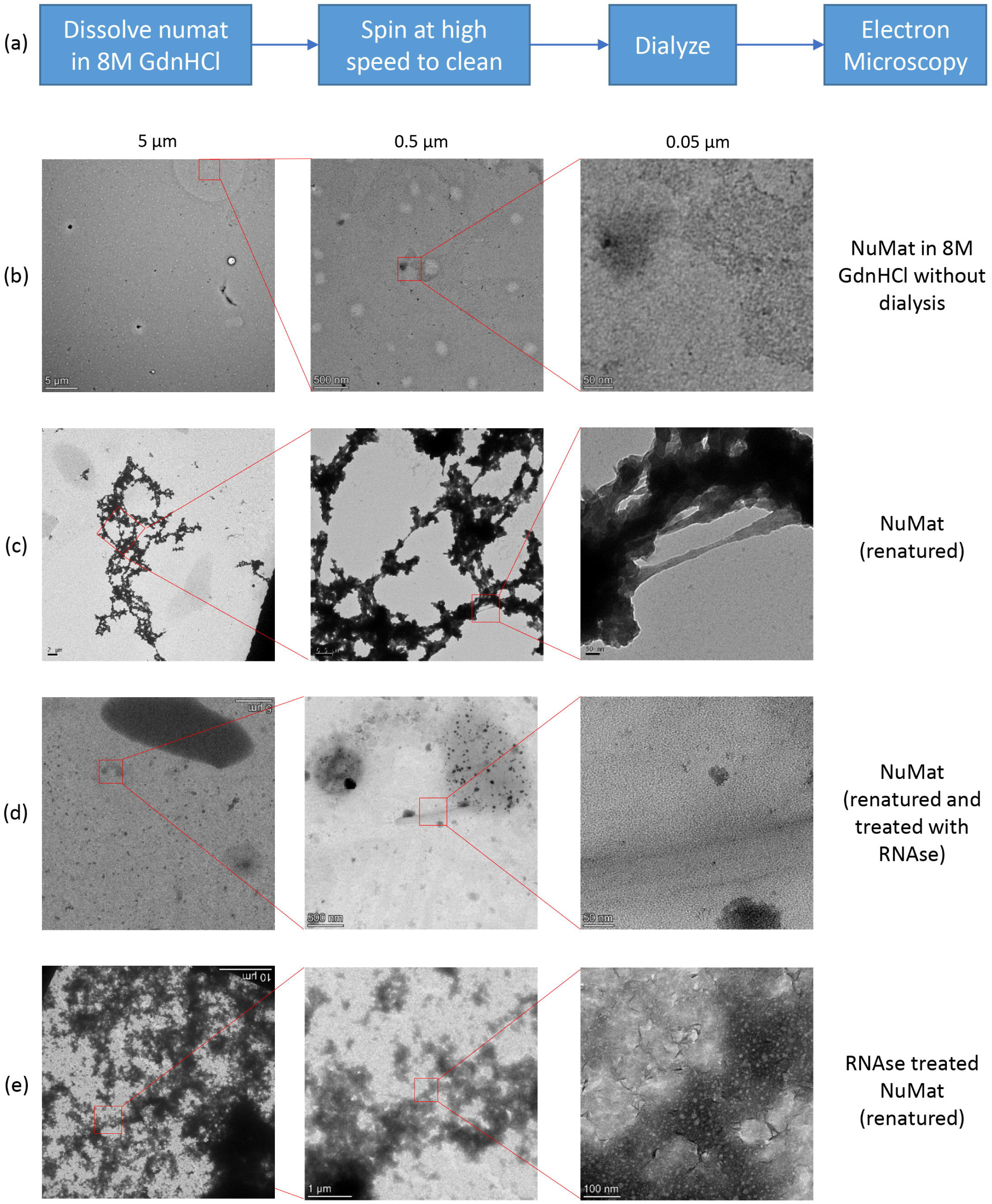
NuMat proteins self-assemble in a network of fibers. NuMat proteins were dissolved in 8M GdnHCl and any debris was removed by high-speed spin, schematic shown in **(a)**. The lack of ultrastructure in the 8M denaturant is shown in **(b)**. This solution, when dialyzed to remove GdnHCl, forms networks of fibers (shown in **(c)**). When the fibers obtained in (c) were treated with RNAse, the fibers collapsed (shown in **(d)**). Similarly, NuMat preparation with RNAse treatment did not yield a contiguous fiber upon renaturation (shown in (e)). For all experiments, pictures were taken at three different magnifications (scales of 5 µm, 0.5 µm, and 0.05 µm unless specified). The magnified area is outlined in a red square.

The phenomenon of fiber formation by self-association is not observed in the renaturation of BSA or nuclear proteins extracted during NuMat preparation at DNase I digestion step, and salt extraction step. This implies that not all sets of proteins can assemble through homotypic or heterotypic self-association. Thus, NuMat consists of proteins that have the propensity to be assembled in this manner (Fig EV10 a, b, and c). To further understand the relationship of protein solubility with the propensity for reassembly through renaturation, we dialyzed out GdnHCl from the extraction fractions. The presence of renatured fibers was seen only in the 8M fraction (Fig 4). Thus, the RNA-dependent homotypic or heterotypic self-association of difficult-to-solubilize NuMat proteins results in the formation of these fibers. Furthermore, it is likely that these findings would apply to the completely insoluble fraction (P) (Fig 1) obtained through sequential extraction, left out in the renaturation experiments because of its insolubility.

**Figure 4:**
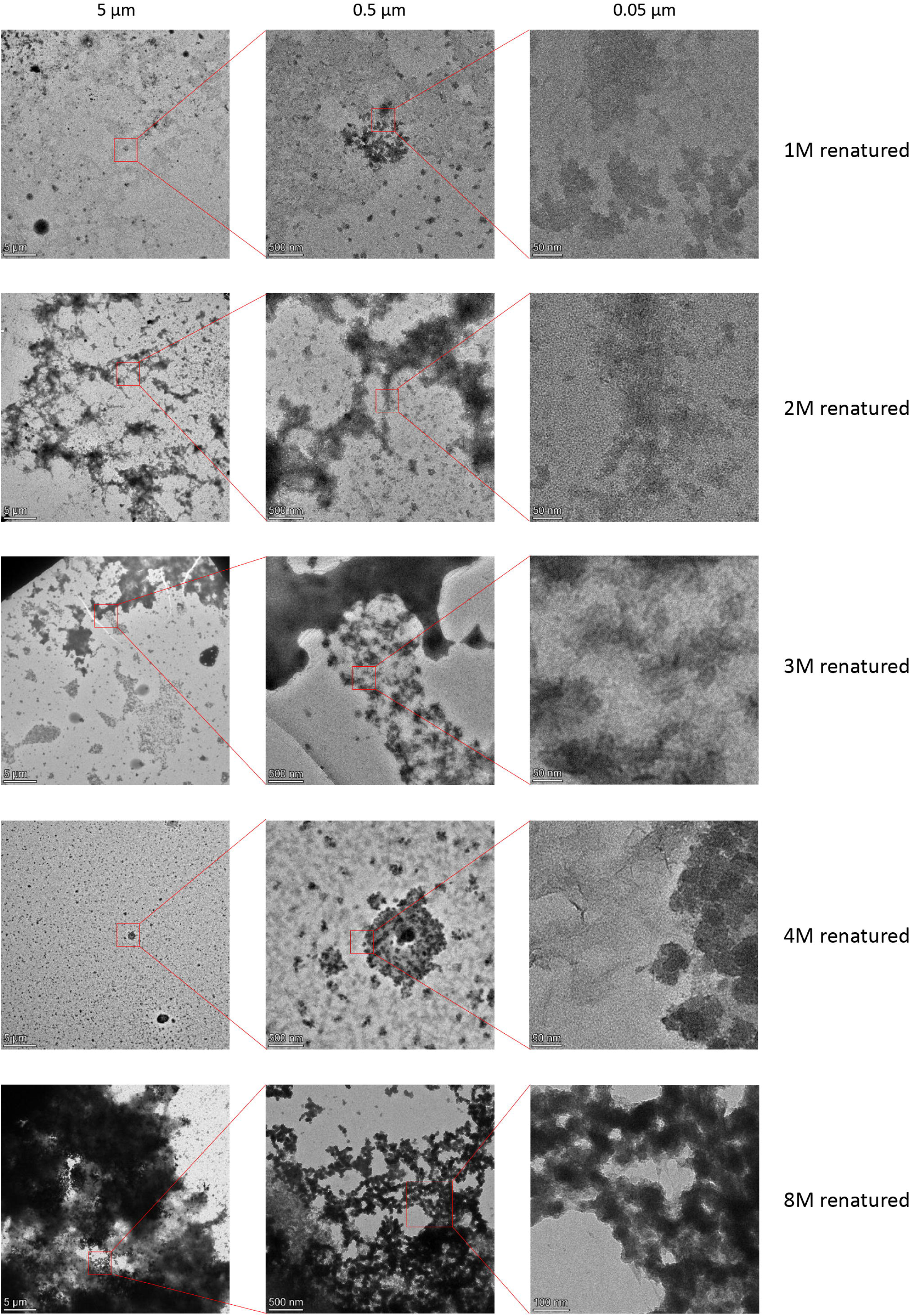
Difficultly-solubilized proteins contribute the most to self-assembling fibers. The extraction fractions of NuMat prepared from sequential extraction by GdnHCl (schematic in Fig 1 a) were renatured (schematic in Fig 3 a), spun, resuspended in 10 µL water and spotted on the grid. Each row shows electron micrographs for each extraction fraction renaturate (1M renatured, 2M renatured, 3M renatured, 4M renatured, and 8M renatured, as labelled) at three different magnifications (at scales 5 µm, 0.5 µm, and 0.05 µm, marked on top of each column). The area magnified from one magnification to the next is shown using a red box.

### Proteomics of renatured fraction

To understand further the nature of the NuMat proteins renaturing through self-association, we pelleted down the renatured fibers from the dialyzed suspension and performed proteomics on them. Principal Component Analysis of this dataset with extraction fractions and non-fiber forming fractions (DNase I digestion fraction and Salt Extraction fraction) shows that this set of proteins is very distinct from non-fiber forming fractions (Fig 5 a). Comparison of fiber-forming renaturation fraction with non-fiber forming fractions yields a cluster of 198 proteins – with statistically significant and high fold change enrichment in renaturation fraction (Fig 5 b, and c) (Supplementary Table 7 sheet 1), which we consider as core renaturation components. These factors are enriched in GO terms related to histone modification, transcription, and chromosome organization (Fig 5 d, Supplementary Table 8 sheet 1). When comparing this cluster with individual extraction fractions, we find that the largest overlap is with the 8M fraction, agreeing with the observations from fraction-wise renaturation (Fig 4 e, Fig 5 e, Supplementary Table 7, sheet 2). Here, we see that the renaturation overlap proteins only present in 8M and absent in other extraction fractions are GO enriched with terms related to transcription regulation, and chromosome organization (Fig EV11 a, Supplementary Table 8 sheet 2). In contrast, renaturation overlap proteins common in all fractions are enriched in terms related to translation and ribosomal biogenesis (Fig EV11 b, Supplementary Table 8 sheet 2). The abundance of RNA-binding proteins in renaturing factors reasserts the importance of RNA in bringing large complexes together and setting a structural framework. This explains why the reassembly collapses when RNA is eliminated (Fig 3 d).

**Figure 5:**
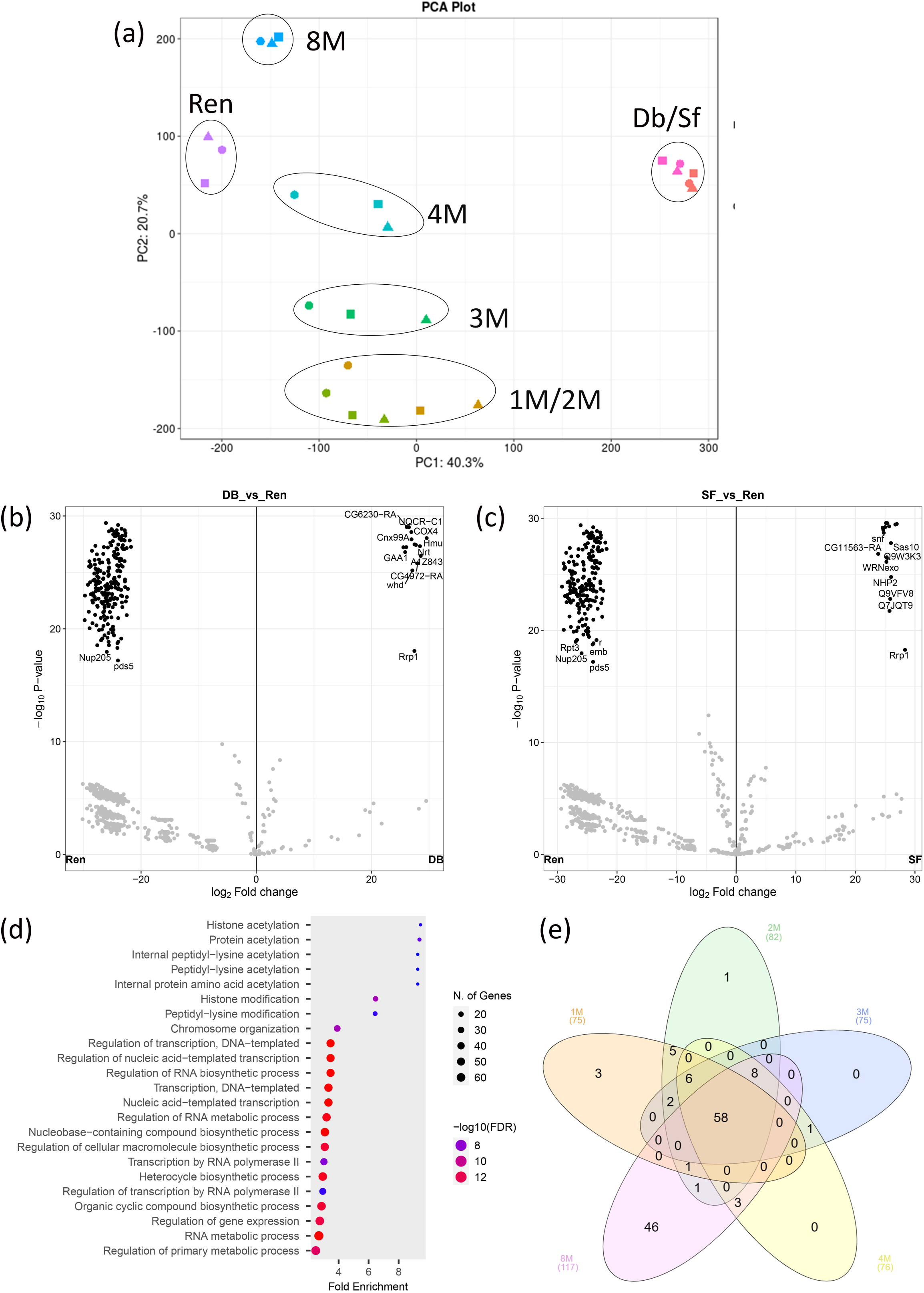
NuMat proteins responsible for reassembly are difficult to solubilize and belong to diverse pathways. **(a)** The proteome profile of renatured pellet is different from non-NuMat nuclear fractions (DNase I digestion fraction and Salt extraction fraction) and similar to the 8M extraction fraction. Shown here is the PCA plot where the replicates of individual fractions cluster together. The renatured fiber-forming fraction clusters farthest away from non-fiber forming fractions (DNAse I digestion fraction and salt extraction fraction), and closer to the extraction fractions, especially 8M. **(b)** Quantitative comparison of the proteomic profile of renatured fraction with DNase I digestion fraction. The horizontal axis shows the log_2_ fold change of the average LFQ intensity ratio for an individual protein for DNAse I digestion (Db) vs Renaturation (ren), and the vertical axis shows the statistical significance of the fold change in terms of –log10 of the p-value. We see a distinct cluster of 273 proteins that have >2^20^ fold change and <10^-10^ p-value, denoting proteins that are principal exclusive contributors to fiber-forming renaturation fraction (highlighted in red square). **(c)** Quantitative comparison of the proteomic profile of renatured fraction with Salt Extraction fraction. The horizontal axis shows the log_2_ fold change of the average LFQ intensity ratio for an individual protein for Salt Extraction (Sf) vs Renaturation (ren), and the vertical axis shows the statistical significance of the fold change in terms of –log10 of the p-value. We see a distinct cluster of 219 proteins that have >2^20^ fold change and <10^-10^ p-value, denoting proteins that are principal exclusive contributors to fiber-forming renaturation fraction (highlighted in red square). **(d)** Gene Ontology analysis of core renaturation factors (made with ShinyGO) shows high fold change enrichment of GO terms related to biological processes like translation, histone modification, and chromosome organization. Y-axis shows various biological processes enriched in core renaturation factors, X-axis shows fold change, the dot size shows the absolute protein count for a pathway, and the dot color represents the p-value of enrichment (red means more significant) **(e)** Contribution of extraction fractions to core renaturation factors. The proteins that from the overlap of core renaturation fraction with individual extraction fractions were used as input in a 5-way Venn diagram (made with interactivenn). We see that the biggest contribution to CRF comes from 8M, both unique and overlapping.

## DISCUSSION

NuMat is the insoluble component of the eukaryotic cell nucleus, resistant to dissolution upon DNase I and salt-detergent treatment, and retains the spatial organization of the nucleus (Berezney and Coffey, 1974). In absence of characterized subfractions of NuMat, we establish GdnHCl-based stepwise extraction as a method to gradually and exhaustively solubilize NuMat proteins.

Several investigations on the whole cell proteome have identified highly insoluble proteins in mammalian cells. A small fraction of proteins change their solubility status upon binding ATP and RNA – and are implicated in the formation of condensate scaffolds through PTMs and signalling (Sridharan et al., 2019, 2022). However, a large number of insoluble proteins do not change their solubility status. Using stepwise extraction with different concentrations of GdnHCl provides a way to further dissect and analyze the difficult-to-solubilize proteome (Mateus et al., 2021) – which we do here for the nuclear non-chromatin insoluble fraction, *i.e.*, NuMat. Given the recent emergence of observations showing semi-solid and solid condensates in the subnuclear space, the solubility profile of NuMat can be utilized to further understand the structural basis of nuclear compartmentalization.

While both GdnHCl and Urea are chaotropes able to unfold the proteins, GdnHCl can shield ionic interactions due to its charge (Biswas et al., 2018). Therefore, the observation that different concentrations of GdnHCl but not of Urea can solubilize proteins from NuMat indicates that the association of the solubilized protein with the undissolved substratum is ionic (Fig EV2 a, and b). To check if the partial solubilization of NuMat is due to the saturation of the extraction buffer with proteins, we analyze the pattern of protein solubilization with multiple sequential extractions of NuMat at the same concentration of GdnHCl. We observe that all GdnHCl extraction concentrations solubilize a finite amount of protein in a finite amount of time, with negligible proteinaceous material leeching upon further extraction (Fig EV3). Thus, individual extraction fractions performed in a sequential manner can be considered distinct from each other (Fig 1 c). The reproducible proteomic profile of GdnHCl-based sequential extraction fractions of NuMat leads us to reason that the sequential solubilization of NuMat proteins is not random and is due to their biophysical propensity to be solubilized at a particular concentration and detach from the undissolved substratum in NuMat (Fig 1 c). This could either be due to biophysical properties intrinsic to the proteins’ amino acid sequence or the creation of various biophysical subspecies through PTMs and heterotypic or homotypic macromolecular interactions. Upon analysis of the proteomic profiles of GdnHCl extractions fractions, we find that the broad biophysical properties of proteins intrinsic to their amino acid sequences that could affect the solubilization behavior pattern of proteins, or where an extraction bias would be reflected, are uniformly distributed across the GdnHCl extraction fractions (Fig EV6). Therefore, a protein’s differential strength of association with the undissolved substratum depends on PTM modifications and macromolecular interactions. The scaffold formation ability of proteins through polymerization or phase separation and their spatial organization has been observed to be modulated by PTMs in the case of Hp1 and Lamin, among others (Naetar et al., 2021; Williams et al., 2020).

Depending on the environment of the protein in various nuclear subcompartments, the organization of the biophysical subspecies of a protein in the subnuclear space extrinsically shifts the protein’s propensity towards being easily solubilized, difficultly solubilized or being subdivided into different subpopulations with different solubilities *i.e.*, ubiquitously-solubilized. Thus, sequential extraction with increasing GdnHCl concentrations clears off loosely bound and accessible material, leaving behind tightly packaged proteins. This is clearly observed in the case of relatively well-characterized complexes like Nuclear Lamina and Nuclear Pore Complex (Maimon et al., 2012; Sapra et al., 2020). In the case of the Lamina, the core structural protein Lamin is present in all fractions but, as seen in the microscopy images (Fig 2, Fig EV7), the retention of the peripheral Lamin mesh is very prominent. This is possibly due to multiple subpopulations of Lamin, some engaged in forming the peripheral mesh, others interacting with various complexes across the nuclear space (Naetar et al., 2021; Parnaik, 2008). Other Nuclear Lamina proteins Man1 and LBR have previously been observed to be unsolubilized by 4M Urea (Dreger et al., 2001) and 8M Urea (Lin et al., 2000). We observe that Man1 is indeed difficult to solubilize (exclusive to 8M). In contrast, Hp1 and LBR are easier to solubilize (depleted in 8M-P), as they anchor membrane and chromatin onto the Lamina (Makatsori et al., 2004). In the case of the Nuclear Pore Complex, the members of the tightly packed pore (membrane anchors (NDC1, Gp210), inner ring complex (Nup93, Nup205, Nup154), outer ring complex (Nup107, Nup133, Nup160, Nup37, Nup44a, Nup75, but not Nup98-96 and sec13), and filamentous mesh on the nuclear side (Tpr or Megator) and cytoplasmic sides (Gle1, mbo, Nup358) are very difficult to solubilize. In contrast, the pore interior proteins (Nup62, Nup58, Nup54) are readily solubilized and are strongly depleted in 8M/P.

Using *in situ* NuMat preparation, we verify that the low-concentration extractions do not disrupt the nuclear shape as inferred from staining for Lamin, wherein a similar trend of extraction at 1M, 2M, and 3M GdnHCl is seen for key NuMat proteins (Fig 2). Additionally, we see that while GAF and RNAP2 are organized in pan-nuclear foci, other NuMat proteins are specific to subnuclear domains, which agrees with the current understanding of nuclear sub-compartmentalization (Zidovska, 2020). This aligns with the idea that the distribution of proteins in NuMat is a snapshot of their distribution in nuclei (Razin et al., 2014). Thus, we are capturing and dissecting the spatial organization of various nuclear substructures together and not the randomly clumped together interchromatin RNA transport channels (Razin and Gromova, 1995). Our observations suggest that NuMat does not behave like a random aggregate in which proteins would have different solubility profiles in different NuMat preps. It also does not behave like a canonical polymeric activity-driven fibrous structure because the unsolubilized fraction retains several key organizational and functional factors (Tpr, Top1/2, SMC2/4, RpII140, RpII215, Prp8, *etc.*). In contrast, fibrous proteins like actin, emerin, myosin, etc., are easily solubilized. It is interesting that tubulins, some kinesin subunits, mini spindles, and α/β-spectrins, are sequestered in the nucleus and are enriched in the 8M/P fraction. These proteins have been implicated in fiber formation and the mechanobiology of nuclear positioning (Davidson et al., 2020) but their possible fiber dynamics in the nuclear interior are yet to be understood.

It should be noted that several recent studies have implicated RNA-protein interactions and subsequent condensate formation in the self-assembly of nuclear subcompartment scaffolds. These scaffolds have been seen to behave in liquid, solid, and semi-solid manners and cause the enrichment of different subspecies of subcompartment-associated markers (Lafontaine et al., 2021; Musacchio, 2022). Therefore, we explored the possibility of the *in vitro* reassembly of NuMat components. To check if NuMat proteins can independently self-assemble, we dismantle the NuMat in 8M GdnHCl and ask if it can renature and form ultrastructural features if the chaotrope is removed by dialysis. As shown in Fig 3, NuMat proteins solubilized by 8M GdnHCl show a unique propensity to self-assemble into fibers. No reassembly is seen upon renaturing protein suspension of similar concentration consisting of chromatin-associated proteins extracted in DNase I/salt fraction (obtained during NuMat preparation) (Fig EV10 a, and b). Upon comparing the proteome of the renatured NuMat fibers to the proteome of DNase I and salt extraction fractions, a cluster of proteins enriched in renatured fibers emerges (Fig 5 b, and c). We reason that these proteins are highly likely to be causative to fiber formation through heterotypic or homotypic association among themselves, mediated by RNA (Fig 3 c, and d). This further emphasizes the role of RNA in the architecture of the nuclear interior as claimed in old electron microscopic studies of NuMat fibers (He et al., 1990) and recent observation of key NuMat proteins (MATR3, NUMA, SAF-A) forming a subnuclear RNP network segregated from chromatin (Creamer et al., 2021).

The renaturation of extraction fractions combined with the analysis of the overlap of the core renaturation components with extraction fractions shows that the difficult-to-solubilize subpopulations of endogenous nuclear proteins readily renature (Fig 4 e, and Fig 5 e). The component of renaturation factors that is unique to 8M extraction fraction is enriched in GO terms related to chromosome organization, histone modifiers and control of transcription (Fig EV11 a). In contrast, the more ubiquitously-solubilized protein component is enriched in GO terms associated with ribosomal subunit precursors and splicing (Fig EV11 b), showing functional divergence between the two subtypes. We speculate that the difficultly-solubilized proteins form domains with RNA acting as an indispensable building block of the pan-nuclear architectural framework.

In summary, we strip NuMat down sequentially, based on the differential solubility of its constituents, dissecting the spatial organization of proteins in the non-chromatin nuclear organization. The NuMat proteins form a multi-layered organization through RNA-dependent mutual association. As we observe mosaicism in the protein components and the architectural importance of RNA in the reassembly of NuMat, we suggest that instead of canonical homopolymeric fibers, a non-canonical multicomponent assembly is involved in fibrous meshwork formation. Additionally, the solubility profile of this meshwork provides a proteome-wide biophysical tool to examine the dynamics of insoluble endogenous nuclear proteins and the liquid/solid/semi-solid scaffolds in various nuclear substructures they engage with. Whether the heterogeneity inherently embedded in such a structure brings in the necessary functional multiplicity remains to be explored. The *in situ* NuMat preparation combined with *Drosophila* genetics and cell biology approaches, will accelerate the exploration of key NuMat components outlined here in the ultrastructural context.

## EXPERIMENTAL PROCEDURES

### A: Preparation of NuMat from *Drosophila melanogaster* embryos

The protocol for the preparation of NuMat has previously been described in (Sureka and Mishra, 2021) and used here with minor modifications, described as follows.

Embryos from Canton S flies were collected overnight (16 h), and dechorionated in 50% bleach (*i.e.*, 2% Sodium Hypochlorite). The dechorionated embryos were lysed in freshly prepared Nuclear Isolation Buffer I (NIB I: 10 mM Tris-Cl pH 7.4, 40 mM KCl, 1 mM EDTA, 0.1 mM EGTA, 0.25 mM Spermidine, 0.1 mM Spermine, 0.1 mM PMSF, 0.25 M Sucrose) with Potter-Elvehjem-Type tissue grinders (6 down-strokes and 6 up-strokes). The homogenized lysate was filtered with Miracloth of pore size 22-25 µm and the filtered lysate was spun at 600g at 4°C for 1 min to remove debris. The crude nuclear suspension supernatant thus obtained was layered over an equal volume of Nuclear Isolation Buffer II (NIB II: 10 mM Tris-Cl pH 7.4, 40 mM KCl, 1 mM EDTA, 0.1 mM EGTA, 0.25 mM Spermidine, 0.1 mM Spermine, 0.1 mM PMSF, 1.8 M Sucrose) in an Oak Ridge style tube and spun at 6000g (4°C, 10 minutes). The pellet containing nuclei was washed thrice with NIB I at 1000g, 4°C for 10 minutes. The quality check for nuclear purity was done by microscopy (DAPI staining, and DIC) during the experiment and western blot (probed for Lamin Dm0 and HisH3) afterward; images of western blot are shown in Fig EV1 a. SEM images of purified nuclei are shown in EV1 b.

Purified nuclei were stabilized in NIB I at 37°C for 20 minutes. At this step, we also lysed 2 µL of the nuclear suspension in 0.5% SDS and measured optical density units (ODU) at 260 nm to quantify nuclei in the suspension by quantifying total DNA. The stabilized nuclei were pelleted (at 1000g, 4°C for 10 minutes) and resuspended in DNase I digestion buffer (20 mM Tris-Cl pH 7.4, 20 mM KCl, 70 mM NaCl, 10 mM MgCl_2_, 0.5% Triton X-100, µg/mL DNase I, 0.1 mM PMSF; DNase I added after resuspension). The suspension was incubated at 4°C for 1 hour. Nuclei were pelleted at 1000g at 4°C for 10 minutes. The pellet was resuspended in extraction buffer I (5 mM HEPES pH 7.4, 2 mM KCl, 2 mM EDTA, 0.25 mM Spermidine, 0.1 mM Spermine, 0.1 mM PMSF, 0.5% Triton X-100, 0.4 M NaCl) and in extraction buffer II (5 mM HEPES pH 7.4, 2 mM KCl, 2 mM EDTA, 0.25 mM Spermidine, 0.1 mM Spermine, 0.1 mM PMSF, 0.5% Triton X-100, 2 M NaCl) sequentially and rotated (10 rpm, RT for 10 minutes in each case). The suspension was pelleted at 3000g for 15 minutes at RT. To keep track of quantity, NuMat preparation was resuspended in extraction buffer II (10 µL per ODU) and the total suspension was split into 500 µL or 50 ODU equivalents. These suspensions were spun down, washed twice in washing buffer (WB: 10 mM Tris-Cl pH 7.4, 20 mM KCl, 1 mM EDTA, 0.25 mM Spermidine, 0.1 mM Spermine, 0.1 mM PMSF) and were stored at −30°C. A silver-stained gel profile of NuMat proteins in contrast with an equivalent amount of nuclei, DNase I digestion supernatant, and salt extraction supernatant is shown in EV1 c. Retention of architectural protein Lamin Dm0 and removal of chromatin protein Histone H3 is shown in western blot in EV1 d.

### B: Subfractionation of NuMat with Guanidinium Hydrochloride (GdnHCl)

NuMat was subfractionated by sequential extraction with GdnHCl extraction buffers (1M, 2M, 3M, 4M, 8M) as per the experimental schematic shown in Fig 1 a. The experimental conditions mentioned in Fig 1 a were finalized through the standardization process described in Supplementary file 1.

The NuMat aliquot (derived from 50 ODU nuclei) was resuspended in 200 µL 1M GdnHCl (in 20 mM MOPS, pH 7.4), extracted for 2 hours at room temperature, and pelleted down (5000g at RT for 5 min). The supernatant was removed and saved for proteomics at −30°C. The pellet from the extraction was resuspended in 200 µL GdnHCl buffer of the next concentration for the next extraction. The sequential extraction process subfractionates the NuMat preparation in 6 parts: 1M extract, 2M extract, 3M extract, 4M extract, 8M extract, and Pellet – encircled in red in Fig 1 a.

### C: SDS-PAGE of GdnHCl fractions

To ensure a good SDS-PAGE run and proper size resolution, all the samples loaded on SDS-PAGE have to have equal GdnHCl concentrations. For this purpose, 80 µL extracts from each GdnHCl fraction (20 ODU equivalent) were taken in separate tubes. A 20 µL mixture of 8M GdnHCl and Milli-Q water prepared in different proportions for individual fractions was added to the respective extracts to adjust the concentration to 3M. The 100 µL extracts thus obtained were boiled with 25 µL 5x laemmli buffer and loaded on the gel. The gel was run at 80 volts till the time the GdnHCl aggregate formed in the wells and the dye front entered the stacking gel. At this point, the wells were flushed and the buffer in the gel compartment was changed. After this, the gel was run like a regular SDS-PAGE gel.

### D: Renaturation

The experimental schematic for renaturation is shown in Fig 3 a. The NuMat aliquot or the other preparations as specified in Fig 3 were dissolved in 200 µL 8M GdnHCl by vigorous vortexing. The solution was pre-cleared of debris by centrifugation at 21000g for 10 min at RT. The supernatant was dialyzed against 20 mM MOPS buffer (pH 7.4) overnight. A carbon-coated copper/nickel grid was dipped in the dialyzed renaturate and shaken lightly. The liquid was removed using a blotting sheet and the grid was stained with uranyl acetate. The samples were visualized in Jeol JEM-2100 (200 kV) or Thermo Fisher Talos L120C (120 kV) Transmission Electron Microscopes. For proteomics, the NuMat renaturate was spun down in a pellet and processed.

### E: Proteomics and Data Analysis

GdnHCl SDS-PAGE gels of the samples were Coomassie stained. Each lane was cut into three strips of higher, medium, and low molecular weight strips. Each strip was separately diced into 1 mm^3^ pieces which were washed three times with wash buffer (25 mM NH_4_HCO_3_, 50% Acetonitrile). The pieces were dehydrated using 100% Acetonitrile and dried in a SpeedVac followed by 10 mM DTT treatment in 25 mM NH_4_HCO_3_ (45 minutes at 55°C). After this, the cubes were saturated with 55 mM Iodoacetamide (30 minutes at 37°C). The pieces were dried again with 100% Acetonitrile and subsequent SpeedVac. The dried pieces were wetted in 10 ng/µL trypsin gold (Promega) in 25 mM NH_4_HCO_3_ and incubated at 37°C overnight (16 h). After 16 h, trypsin action was stopped and peptides were eluted with 2.5% TFA in 50% Acetonitrile in two sequential washes. The supernatant from both washes was collected in fresh tubes and dried with SpeedVac. The pellet was resuspended in 0.1% TFA and 2% Acetonitrile. This sample was desalted with Millipore p10 Ziptip C18. The elution buffer was evaporated using SpeedVac and the peptides were resuspended in 0.1% formic acid and 2% Acetonitrile.

The spectra were collected on Thermo Scientific Q-Exactive HF on a 60-minute acetonitrile gradient and analyzed with MaxQuant (version 1.5.7.4) at default settings. Search parameters of this setting are provided in Supplementary Table 1. Proteins that had at least two unique peptides and non-zero Label Free Quantitation (LFQ) intensities across all three biological replicates were considered for the analysis. All the replicates of all the fractions were subjected to Principal Component Analysis (PCA), considering the correlation between LFQ intensities associated with all the protein lists of individual samples (Shah et al., 2019). The differential enrichment analysis for these proteins’ LFQ intensities across different fractions was performed with DEP and LFQ-analyst (Shah et al., 2019; Zhang et al., 2018). Proteins that showed more than 25log_2_ fold LFQ intensity difference statistically significantly (p<0.001) were chosen. Their differential enrichment score was plotted in a heatmap using ClustVis, with average distance clustering of rows (Metsalu and Vilo, 2015). Interactivenn and ShinyGO have been used to look at the distribution and pathway enrichment of protein lists (Ge et al., 2020; Heberle et al., 2015).

### F. *In situ* NuMat preparation and GdnHCl extraction

The protocol for preparing *in situ* NuMat has previously been published (Pathak et al., 2022a, 2022b). To mimic the extraction procedure described in methods section B, we eliminated formaldehyde crosslinking. The modified procedure and subsequent sequential extractions are described as follows.

Embryos collected from Canton S flies were washed in 50% bleach until dechorionated. For the removal of the vitelline membrane, the embryos were rotated with 50% heptane in 1X PBS for 10 minutes. The embryos at the heptane-PBS interface were taken for further processing in a heptane-methanol (1:1) mix. After this step, the embryos that sat at the bottom of the tube were washed with cold methanol. After the fixed embryos became non-sticky, they were stored in methanol at −30°C. For NuMat preparation, embryos were rehydrated by drop-by-drop addition of PBT (1X PBS, pH 7.4 with 0.5% Triton X-100) and washed thrice with cold PBT (20 minutes each time). Next, the embryos were stabilized at 37°C for 20 minutes. After this, the embryos were treated with Extraction Buffer I (0.4 M NaCl) and Extraction buffer II (2 M NaCl), for 20 minutes each. The stabilized embryos were submerged in DNase I digestion buffer (30 minutes). The resulting *in situ* NuMat was washed thrice with cold PBT (20 minutes each time).

For sequential extraction of *in situ* NuMat, 1M GdnHCl solution was added to three aliquots of *in situ* NuMat and rotated at room temperature for 2 hours – the solution was carefully aspirated at the end. One aliquot was subjected to three PBT washes, 20 minutes each. Meanwhile, 2M GdnHCl solution was added to the other two aliquots and rotated at room temperature for 2 hours. One aliquot was subjected to three PBT washes, 20 minutes each. Meanwhile, 3M GdnHCl solution was added to the remaining aliquot and rotated at room temperature for 2 hours. This aliquot was subjected to three PBT washes, 20 minutes each. The embryo structure collapses at GdnHCl concentrations higher than 3M.

### G. Immunostaining and confocal imaging of extracted *in situ* NuMat

The embryo derivative samples (Embryos, *in situ* NuMat, and *in situ* NuMat extracted 1M, 2M, 3M GdnHCl buffers) were blocked by rotating in 3% BSA in PBT for 2 hours at 4°C. After a quick rinse with cold PBT, primary antibody solution was added and the suspension was rotated overnight at 4°C (16 h). Next, after three washes with cold PBT, the sample was rotated with secondary antibody solution (4°C for 3 h). The antibody solution was aspirated and DAPI was added in appropriate amounts to the sample bed volume. The sample was washed thrice with cold PBT, 20 minutes each time. The sample was mounted in Vectashield mounting media without DAPI and sealed.

Mounted samples were imaged with Olympus FV3000 confocal microscope at 100x magnification with 3x zoom, taking near-saturation voltage values. The raw data were processed with ImageJ to get the same number of slices of all scans of one sample, to take the sum intensity projection, and to add the scale bar.

Details of important reagents and antibodies are specified in tables in Supplementary file 1.

### H. Experimental Design and Statistical Rationale

We have taken two GdnHCl-based approaches to chisel at NuMat: first, to subfractionate it with finite extractions with increasing concentrations of GdnHCl, and second, to dissolve it in GdnHCl and remove it through dialysis to check for reassembly. For extraction subfractionation, the profiles of the proteins extracted in different fractions are deciphered by LC-MS-MS-based proteomics. For both lines of investigation, three biological replicates of each fraction were analyzed. The SDS-PAGE lane of each replicate was cut into three strips based on size to improve proteomics coverage and their data were merged in the MaxQuant run by specifying the spectra from strips as fractions of the same replicate.

Only proteins identified in all three biological replicates with two or more peptides and non-zero total LFQ intensity were considered true fraction components and were used for further analysis. Principal Component Analysis (with LFQ-Analyst) was used to judge the reliability of quantification measurements between biological replicates. The differential enrichment of protein across GdnHCl extraction fractions was judged by the fold change difference (>25log_2_-fold) between triplicate LFQ-intensities and the associated p-value (<0.001) (using LFQ-analyst and DEP). For identifying the proteins involved in reassembly, we compared NuMat renaturation fraction with DNase I digestion fraction and salt extraction fraction, we have looked at the cluster emerging through the volcano plot and subsequent GO analysis of the subset of this cluster, which does show very high fold change (>20log_2_-fold) and p-value (<0.001).

## AUTHOR CONTRIBUTIONS

R.K.M. conceptualized and R.K.M. and A.B. designed the study. A.B. executed the experiments. A.B. and A.A. cleaned and analyzed the data. R.K.M. and A.B. interpreted the data and wrote the manuscript.

## Supporting information

Extended view figure 1

Extended view figure 2

Extended view figure 3

Extended view figure 4

Extended view figure 5

Extended view figure 6

Extended view figure 7

Extended view figure 8

Extended view figure 9

Extended view figure 10

Extended view figure 11

## ACKNOWLEDGEMENTS

We thank the CCMB community, especially R.K.M. lab members for critical insights and inputs. We thank the Advanced Microscopy and Imaging Facility (C. Subbalaxmi for confocal microscopy and H. Adicherla for electron microscopy) and Proteomics Facility (B Raman) at CCMB.

## DATA AVAILABILITY

The raw proteomics data are available via ProteomeXchange with the identifier PXD031296 (login: **Username**: **Password**: 7FmlIU1M). The filenames corresponding to biological replicates and their fractions have been provided in Supplementary Table 1.

## Supplemental data

This article contains supplemental data.

## Expanded view figure legends

**Expanded view figure 1:** Quality checks of NuMat preparation. (a) The nuclear lysate was run alongside whole cell lysate and probed for Lamin Dm0, GAPDH, and Cytochrome C. While both lanes have the same amount of lamin, the nuclear lysate does not have any GAPDH or Cytochrome C (b) Scanning electron microscope images of the isolated nuclei; there are no cytoplasmic attachments on the outer surface, scale bars in the images. NuMat aliquot alongside nuclear lysate, DNAse I digestion supernatant, and salt extraction supernatant shown in (a) Silver stain of the gel (b) western blot probed for Lamin Dm0 and Histone H3. We see that histones are extracted in salt fraction while the nuclear matrix retains higher molecular weight proteins like Lamin.

**Expanded view figure 2:** Protein profile of the NuMat extraction with GdnHCl and Urea. Four separate NuMat aliquots were resuspended in 1M, 2M, 3M, and 4M GdnHCl solution, and another four in 1.5M, 3M, 4.5M, and 6M Urea. Supernatants were loaded in lanes labelled XM_Sup. Pellets were loaded in lanes labelled as XM_P. Large-scale solubilization is visible in GdnHCl supernatants (a) and none in Urea supernatants (b). There is a marked difference between the proteins solubilized by GdnHCl and the proteins left in the pellet.

**Expanded view figure 3:** Check for leeching in NuMat extraction with GdnHCl by sequential extraction with the same concentration. Four separate NuMat aliquots were resuspended in 1M, 2M, 3M, and 4M GdnHCl solution. The suspension was pelleted every hour and the pellet was resuspended in a fresh solution of the same GdnHCl concentration. Here shown are the silver stains of the 1-hour protein extraction profile for each concentration. Maximum extraction happens within 2 hours in each case.

**Expanded view figure 4:** NuMat aliquot was sequentially extracted with 1M, 2M, 3M, and 4M GdnHCl solutions. All four supernatants and the pellet were loaded. The profiles are slightly different.

**Expanded view figure 5:** Overlap of proteins across fractions, compared by 6-way Venn diagram: We observe that the qualitative profiles of 1M, 2M, 3M, 4M, 8M, and P fractions have 132 proteins common to all fractions (highlighted in a black square). High-concentration extractions (8M and P, highlighted in red circles) and low-concentration extractions (1M and 2M, highlighted in green circles), emerge as larger unique clusters as opposed to 3M-4M. Thus, there is a clear divergence in easily-solubilized (green) and difficultly-solubilized (red) groups.

**Expanded view figure 6:** Comparisons of broad biophysical property histograms across GdnHCl fractions: Shown here are violin plots of protein number distribution for (a) Molecular weight, (b) Isoelectric point, (c) Intrinsic Disorder Percentage (data from MobiDB (Piovesan et al., 2022)), (d) Non-polar Amino Acids for all three replicates of each fraction. There is no biophysical bias in the extraction of embryo NuMat proteins at different GdnHCl concentrations.

**Expanded view figure 7:** Each *in situ* NuMat preparation at each extraction step (Nu for whole nuclei, Nm for in situ NuMat, 1M, 2M, and 3M) was stained with DAPI and immunostained for Lamin Dm0 alongside the target protein ((a) HP1 (b) Pc (c) Trl (d) Tpr/Megator (e) Fib (f) RNAP2. The figures show the sum projection of a few slices from the sheet of nuclei images at 100x magnification and 3x zoom, the Lamin staining showing that the nuclear morphology is intact through the sequential extraction, and DAPI staining showing that the chromatin has properly been removed during *in situ* NuMat preparation. Scale bar 5 µm in all images.

**Expanded view figure 8:** LFQ-intensity-based extraction patterns of key NuMat proteins. Shown here is the extraction pattern of NuMat protein examined through immunofluorescence and confocal microscopy of *in situ* Nuclear Matrix as line diagrams in terms of percentage extraction at each extraction step, X-axis showing the extraction steps (1M to P) and Y-axis showing percentage LFQ-intensity (0-100). Color codes for individual proteins are shown at the side of the graph. We observe that RNAP2 and Megator show no solubilization in the first three steps, HP1 and Pc are solubilized easily while GAF and Fib are ubiquitously-solubilized. Lamin shows easy solubilization here and ubiquitously solubilization in *in situ* Nuclear Matrix.

**Expanded view figure 9:** Non-NuMat nuclear proteins don’t self-assemble to form fibers. (a) DNase I extracted chromatin-associated proteins, (b) Salt extracted nuclear proteins (c) homogeneous protein solution with a single member, BSA. For all experiments, pictures were taken at three different magnifications (scales of 5 µm, 0.5 µm, and 0.05 µm). In the case of extreme magnification from 0.5 µm scale to 50 nm scale, the magnified area is outlined in a red circle.

**Expanded view figure 10:** Morphology of renatured NuMat fibers upon physical perturbations: (a) NuMat renaturation fibers formed by dialysis and the grid dipped in renaturate (b) NuMat renaturation fibers formed by dialysis were spun down at 21000xg, resuspended in 10 µL water and spotted on the grid (c) NuMat renaturation fibers formed by serial dilution (8M->4M->2M->1M->0.5M) and 10 µL renaturate spotted on the grid.

**Expanded view figure 11:** Gene ontology analysis of two major subsets of core renaturation factors, shown here are dot plots where Y-axis shows various biological processes enriched, X-axis shows fold change, the dot size shows absolute protein count for a pathway and the dot color represents the p-value of enrichment (red means more significant). (a) Gene Ontology analysis of core renaturation proteins overlap with proteins unique to 8M (b) Gene ontology analysis of core renaturation proteins overlap with 8M proteins that are common to all fractions. Plotted with ShinyGO.

## REFERENCES

1. Adolph, K.W., Cheng, S.M., Paulson, J.R., Laemmli, U.K., and Brown, J.A. (1977). Isolation of a protein scaffold from mitotic HeLa cell chromosomes. Biochemistry 74, 4937–4941.

2. Agutter, P.S., and Richardson, J.C.W. (1980). Nuclear non-chromatin proteinaceous structures: Their role in the organization and function of the interphase nucleus. J. Cell Sci. *VOL.*44, 395– 435.

3. Albiez, H., Cremer, M., Tiberi, C., Vecchio, L., Schermelleh, L., Dittrich, S., Küpper, K., Joffe, B., Thormeyer, T., Von Hase, J., et al. (2006). Chromatin domains and the interchromatin compartment form structurally defined and functionally interacting nuclear networks. Chromosom. Res.

4. Barutcu, A.R., Blencowe, B.J., and Rinn, J.L. (2019). Differential contribution of steady state RNA and active transcription in chromatin organization. EMBO Rep. 20, e48068.

5. Berezney, R., and Coffey, D.S. (1974). Identification of a nuclear protein matrix. Biochem. Biophys. Res. Commun. 60, 1410–1417.

6. Berezney, R., and Coffey, D.S. (1975). Nuclear Protein Matrix: Association with newly synthesized DNA. Science (80-.). 189, 291–293.

7. Berezney, R., Mortillaro, M.J., Ma, H., Wei, X., and Samarabandu, J. (1995). The Nuclear Matrix: a structural milieu for genomic function. Int. Rev. Cytol. 162A, 1–65.

8. Biswas, B., Muttathukattil, A.N., Reddy, G., and Singh, P.C. (2018). Contrasting Effects of Guanidinium Chloride and Urea on the Activity and Unfolding of Lysozyme. ACS Omega.

9. Caridi, C.P., D’agostino, C., Ryu, T., Zapotoczny, G., Delabaere, L., Li, X., Khodaverdian, V.Y., Amaral, N., Lin, E., Rau, A.R., et al. (2018). Nuclear F-actin and myosins drive relocalization of heterochromatic breaks. Nature 559, 54–60.

10. Chubb, J.R., Boyle, S., Perry, P., and Bickmore, W.A. (2002). Chromatin Motion Is Constrained by Association with Nuclear Compartments in Human Cells. Curr. Biol. 12, 439–445.

11. Ciejek, E.M., Tsai, M. jer, and O’Malley, B.W. (1983). Actively transcribed genes are associated with the nuclear matrix. Nature 306, 607–609.

12. Comings, D.E., and Okada, T.A. (1976). Nuclear proteins. III. The fibrillar nature of the nuclear matrix. Exp. Cell Res. 103, 341–360.

13. Cook, P.R., Uptain, S.M., Kane, C.M., Chamberlin, M.J., Myers, V.E., Young, R.A., McCracken, S., Cho, H., Maldonado, E., Scully, R., et al. (1999). The organization of replication and transcription. Science 284, 1790–1795.

14. Costa, S., and Shaw, P. (2006). Chromatin organization and cell fate switch respond to positional information in Arabidopsis. Nature 439, 493–496.

15. Creamer, K.M., Kolpa, H.J., and Lawrence, J.B. (2021). Nascent RNA scaffolds contribute to chromosome territory architecture and counter chromatin compaction. Mol. Cell 81, 3509–3525.e5.

16. Dalton, S. (2015). Linking the Cell Cycle to Cell Fate Decisions. Trends Cell Biol. 25, 592–600.

17. Davidson, P.M., Battistella, A., Déjardin, T., Betz, T., Plastino, J., Borghi, N., Cadot, B., and Sykes, C. (2020). Nesprinu accumulates at the front of the nucleus during confined cell migration. EMBO Rep. 21, e49910.

18. Dreger, M., Bengtsson, L., Schöneberg, T., Otto, H., and Hucho, F. (2001). Nuclear envelope proteomics: Novel integral membrane proteins of the inner nuclear membrane. Proc. Natl. Acad. Sci. U. S. A. 98, 11943–11948.

19. Engelke, R., Riede, J., Hegermann, J., Wuerch, A., Eimer, S., Dengjel, J., and Mittler, G. (2014). The Quantitative Nuclear Matrix Proteome as a Biochemical Snapshot of Nuclear Organization. J. Proteome Res. 13, 3940–3956.

20. Fey, E.G., Wan, K.M., and Penman, S. (1984). Epithelial cytoskeletal framework and Nuclear Matrix-intermediate filament scaffold: three-dimensional organization and protein composition. J. Cell Biol. 98, 1973–1984.

21. Ge, S.X., Jung, D., Jung, D., and Yao, R. (2020). ShinyGO: A graphical gene-set enrichment tool for animals and plants. Bioinformatics 36, 2628–2629.

22. Georgiev, G.P. (1967). The Nature and Biosynthesis of Nuclear Ribonucleic Acids. Prog. Nucleic Acid Res. Mol. Biol. 6, 259–351.

23. Georgiev, G.P., and Chentsov, J.S. (1962). On the structural organization of nucleolochromosomal ribonucleoproteins. Exp. Cell Res. 27, 570–572.

24. He, D.C., Nickerson, J.A., and Penman, S. (1990). Core filaments of the Nuclear Matrix. J. Cell Biol. 110, 569–580.

25. Heberle, H., Meirelles, G.V., da Silva, F.R., Telles, G.P., and Minghim, R. (2015). InteractiVenn: a web-based tool for the analysis of sets through Venn diagrams. BMC Bioinformatics 16, 169.

26. Hozák, P., Hassan, A.B., Jackson, D.A., and Cook, P.R. (1993). Visualization of replication factories attached to a nucleoskeleton. Cell 73, 361–373.

27. Kallappagoudar, S., Varma, P., Pathak, R.U., Senthilkumar, R., and Mishra, R.K. (2010). Nuclear matrix proteome analysis of Drosophila melanogaster. Mol. Cell. Proteomics 9, 2005– 2018.

28. Keenen, M.M., Brown, D., Brennan, L.D., Renger, R., Khoo, H., Carlson, C.R., Huang, B., Grill, S.W., Narlikar, G.J., and Redding, S. (2021). HP1 proteins compact DNA into mechanically and positionally stable phase-separated domains. Elife 10, 1–38.

29. Koehler, D.R., and Hanawalt, P.C. (1996). Recruitment of damaged DNA to the Nuclear Matrix in hamster cells following ultraviolet irradiation. Nucleic Acids Res. 24, 2877–2884.

30. Lafontaine, D.L.J. (2019). Birth of Nucleolar Compartments: Phase Separation-Driven Ribosomal RNA Sorting and Processing. Mol. Cell 76, 694–696.

31. Lafontaine, D.L.J., Riback, J.A., Bascetin, R., and Brangwynne, C.P. (2021). The nucleolus as a multiphase liquid condensate. Nat. Rev. Mol. Cell Biol.

32. Lim, W.K., Rösgen, J., and Englander, S.W. (2009). Urea, but not guanidinium, destabilizes proteins by forming hydrogen bonds to the peptide group. Proc. Natl. Acad. Sci. U. S. A. 106, 2595–2600.

33. Lin, F., Blake, D.L., Callebaut, I., Skerjanc, I.S., Holmer, L., McBurney, M.W., Paulin-Levasseur, M., and Worman, H.J. (2000). MAN1, an inner nuclear membrane protein that shares the LEM domain with lamina-associated polypeptide 2 and emerin. J. Biol. Chem. 275, 4840– 4847.

34. Maimon, T., Elad, N., Dahan, I., and Medalia, O. (2012). The human nuclear pore complex as revealed by cryo-electron tomography. Structure 20, 998–1006.

35. Makatsori, D., Kourmouli, N., Polioudaki, H., Shultz, L.D., McLean, K., Theodoropoulos, P.A., Singh, P.B., and Georgatos, S.D. (2004). The inner nuclear membrane protein lamin B receptor forms distinct microdomains and links epigenetically marked chromatin to the nuclear envelope. J. Biol. Chem.

36. Mateus, A., Savitski, M.M., and Piazza, I. (2021). The rise of proteome wide biophysics. Mol. Syst. Biol.

37. Metsalu, T., and Vilo, J. (2015). ClustVis: a web tool for visualizing clustering of multivariate data using Principal Component Analysis and heatmap. Nucleic Acids Res. 43, W566–W570.

38. Misteli, T. (2005). Concepts in nuclear architecture. BioEssays 27, 477–487.

39. Mullenders, L.H.F., van Leeuwen, A. c. kestere. Van, van Zeeland, A.A., and Natarajan, A.T. (1988). Nuclear Matrix associated DNA is preferentially repaired in normal human fibroblasts, exposed to a low dose of ultraviolet light but not in Cockayne’s syndrome fibroblasts. Nucleic Acids Res. 16, 10607–10622.

40. Musacchio, A. (2022). On the role of phase separation in the biogenesis of membraneless compartments. EMBO J.

41. Naetar, N., Georgiou, K., Knapp, C., Bronshtein, I., Zier, E., Fichtinger, P., Dechat, T., Garini, Y., and Foisner, R. (2021). Lap2alpha maintains a mobile and low assembly state of a-type lamins in the nuclear interior. Elife.

42. Nakagawa, S., and Prasanth, K. V. (2011). EXIST with matrix-associated proteins. Trends Cell Biol.

43. Nelson, W.G., Pienta, K.J., Barrack, E.R., and Coffey, D.S. (1986). The role of the nuclear matrix in the organization and function of DNA. Annu. Rev. Biophys. Biophys. Chem. 15, 457– 475.

44. Neri, L.M., Raymond, Y., Giordano, A., Capitani, S., and Martelli, A.M. (1999). Lamin A is part of the internal nucleoskeleton of human erythroleukemia cells. J. Cell. Physiol. 178, 284–295.

45. Nickerson, J.A., Krochmalnic, G., Wan, K.M., and Penman, S. (1989). Chromatin architecture and nuclear RNA. Proc. Natl. Acad. Sci. U. S. A. 86, 177–181.

46. Nickerson, J.A., Krockmalnic, G., Wan, K.M., and Penman, S. (1997). The Nuclear Matrix revealed by eluting chromatin from a cross-linked nucleus. Proc. Natl. Acad. Sci. U. S. A. 94, 4446–4450.

47. Parnaik, V.K. (2008). Role of Nuclear Lamins in Nuclear Organization, Cellular Signaling, and Inherited Diseases. Int. Rev. Cell Mol. Biol. 266, 157–206.

48. Pathak, R.U., Mamillapalli, A., Rangaraj, N., Kumar, R.P., Vasanthi, D., Mishra, K., and Mishra, R.K. (2013). AAGAG repeat RNA is an essential component of Nuclear Matrix in Drosophila. RNA Biol. 10, 564–571.

49. Pathak, R.U., Srinivasan, A., and Mishra, R.K. (2014). Genome-wide mapping of Matrix attachment regions in Drosophila melanogaster. BMC Genomics 15, 1–10.

50. Pathak, R.U., Bihani, A., Sureka, R., Varma, P., and Mishra, R.K. (2022a). In situ Nuclear Matrix preparation in Drosophila melanogaster embryos/tissues and its use in studying the components of nuclear architecture. Nucleus 13, 116–128.

51. Pathak, R.U., Bihani, A., Sureka, R., and Mishra, R.K. (2022b). In situ Nuclear Matrix preparation in Drosophila melanogaster enabling genetic analysis of the nuclear architecture. STAR Protoc. 3.

52. Pederson, T. (2000). Half a century of “the Nuclear Matrix”. Mol. Biol. Cell 11, 799–805.

53. Phair, R.D., and Misteli, T. (2000). High mobility of proteins in the mammalian cell nucleus. Nature 404, 604–609.

54. Piovesan, D., Del Conte, A., Clementel, D., Monzon, A.M., Bevilacqua, M., Aspromonte, M.C., Iserte, J.A., Orti, F.E., Marino-Buslje, C., and Tosatto, S.C.E. (2022). MobiDB: 10 years of intrinsically disordered proteins. Nucleic Acids Res. gkac1065.

55. Raskar, T., Yeow, C., Niebling, S., Kini, R.M., and Hosur, M. V. (2019). X-ray crystallographic analysis of time-dependent binding of guanidine hydrochloride to HEWL: First steps during protein unfolding. Int. J. Biol. Macromol. 122, 903–913.

56. Razin, S. V., and Gromova, I.I. (1995). The channels model of Nuclear Matrix structure. BioEssays 17, 443–450.

57. Razin, S. V., Iarovaia, O. V., and Vassetzky, Y.S. (2014). A requiem to the Nuclear Matrix: from a controversial concept to 3D organization of the nucleus. Chromosoma 123, 217–224.

58. Sapra, K.T., Qin, Z., Dubrovsky-Gaupp, A., Aebi, U., Müller, D.J., Buehler, M.J., and Medalia, O. (2020). Nonlinear mechanics of lamin filaments and the meshwork topology build an emergent nuclear lamina. Nat. Commun. 11.

59. Shah, A.D., Goode, R.J.A., Huang, C., Powell, D.R., and Schittenhelm, R.B. (2019). Lfq-Analyst: An easy-To-use interactive web platform to analyze and visualize label-free proteomics data preprocessed with maxquant. J. Proteome Res. 204–211.

60. Shankar Narayan, K., Steele, W.J., Smetana, K., and Busch, H. (1967). Ultrastructural aspects of the ribonucleoprotein network in nuclei of walker tumour and rat liver. Exp. Cell Res. 46, 65–77.

61. Smetana, K., Steele, W.J., and Busch, H. (1963). A nuclear ribonucleoprotein network (Academic Press).

62. Sridharan, S., Kurzawa, N., Werner, T., Günthner, I., Helm, D., Huber, W., Bantscheff, M., and Savitski, M.M. (2019). Proteome-wide solubility and thermal stability profiling reveals distinct regulatory roles for ATP. Nat. Commun.

63. Sridharan, S., Hernandez-Armendariz, A., Kurzawa, N., Potel, C.M., Memon, D., Beltrao, P., Bantscheff, M., Huber, W., Cuylen-Haering, S., and Savitski, M.M. (2022). Systematic discovery of biomolecular condensate-specific protein phosphorylation. Nat. Chem. Biol. 2022 1810 18, 1104–1114.

64. Sureka, R., and Mishra, R. (2021). Identification of Evolutionarily Conserved Nuclear Matrix Proteins and Their Prokaryotic Origins. J. Proteome Res. 20, 518–530.

65. Sureka, R., Wadhwa, R., Thakur, S.S., Pathak, R.U., and Mishra, R.K. (2018). Comparison of Nuclear Matrix and mitotic chromosome scaffold proteins in drosophila S2 cells-transmission of hallmarks of nuclear organization through mitosis. Mol. Cell. Proteomics 17, 1965–1978.

66. Tan, J.H., Wooley, J.C., and LeStourgeon, W.M. (2000). Nuclear Matrix-like filaments and fibrogranular complexes form through the rearrangement of specific nuclear ribonucleoproteins. Mol. Biol. Cell 11, 1547–1554.

67. Tumbar, T., and Belmont, A.S. (2001). Interphase movements of a DNA chromosome region modulated by VP16 transcriptional activator. Nat. Cell Biol. 3, 134–139.

68. Tumbar, T., Sudlow, G., and Belmont, A.S. (1999). Large-Scale Chromatin Unfolding and Remodeling Induced by VP16 Acidic Activation Domain. J. Cell Biol. 145.

69. Williams, J.F., Surovtsev, I. V., Schreiner, S.M., Nguyen, H., Hu, Y., Mochrie, S.G.J., and King, M.C. (2020). Phase separation enables heterochromatin domains to do mechanical work. BioRxiv.

70. Young, K.G., and Kothary, R. (2005). Spectrin repeat proteins in the nucleus. BioEssays 27, 144–152.

71. Zbarsky, I.B. (1998). On the history of nuclear matrix manifestation. Cell Res. 8, 99–103.

72. Zbarsky, I.B., and Perevoshchikova, K.A. (1948). On some properties of cell nuclear proteins. In Doklady Akad. Nauk SSSR, pp. 77–80.

73. Zeitlin, S., Parent, A., Silverstein, S., and Efstratiadis, A. (1987). Pre-mRNA splicing and the Nuclear Matrix. Mol. Cell. Biol. 7, 111–120.

74. Zhang, X., Smits, A.H., Van Tilburg, G.B.A., Ovaa, H., Huber, W., and Vermeulen, M. (2018). Proteome-wide identification of ubiquitin interactions using UbIA-MS. Nat. Protoc. 13, 530– 550.

75. Zheng, R., Shen, Z., Tripathi, V., Xuan, Z., Freier, S.M., Bennett, C.F., Prasanth, S.G., and Prasanth, K. V. (2010). Polypurine-repeat-containing RNAs: A novel class of long non-coding RNA in mammalian cells. J. Cell Sci.

76. Zidovska, A. (2020). The rich inner life of the cell nucleus: dynamic organization, active flows, and emergent rheology. Biophys. Rev.

