## Supplementary figures and images for "Biochemical deconstruction and reconstruction of Nuclear Matrix reveals the layers of nuclear organization"

### Extended view figure 1

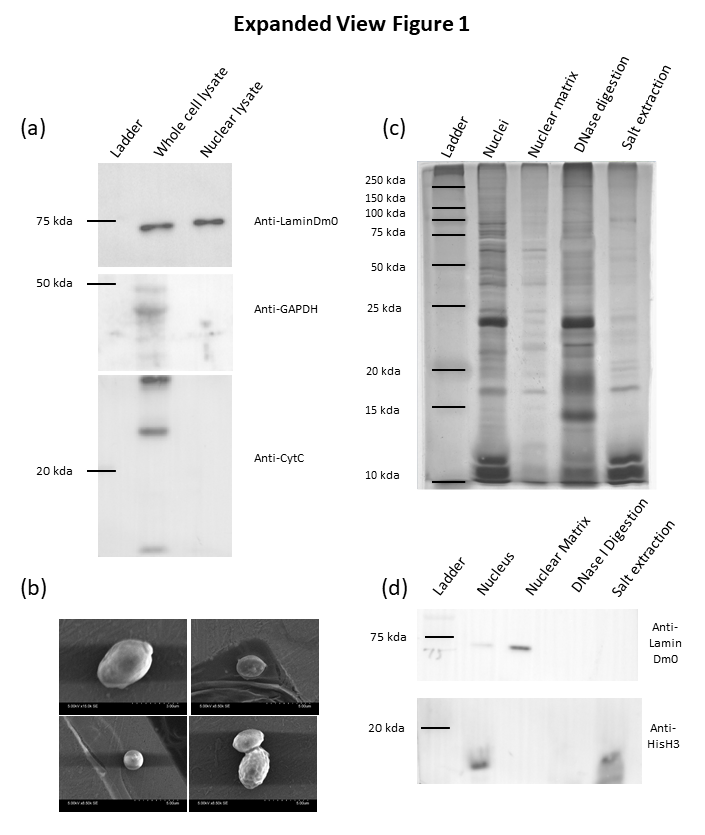

### Extended view figure 2

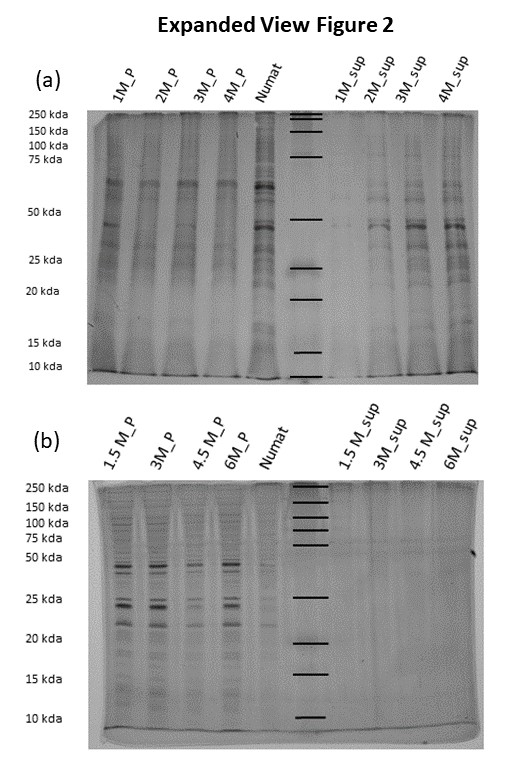

### Extended view figure 3

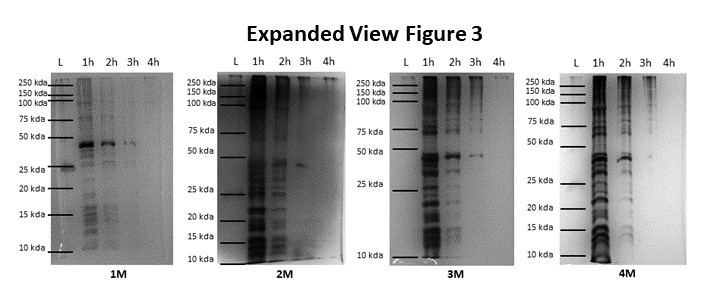

### Extended view figure 4

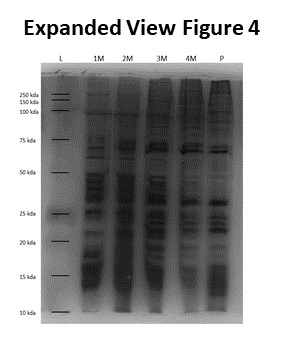

### Extended view figure 5

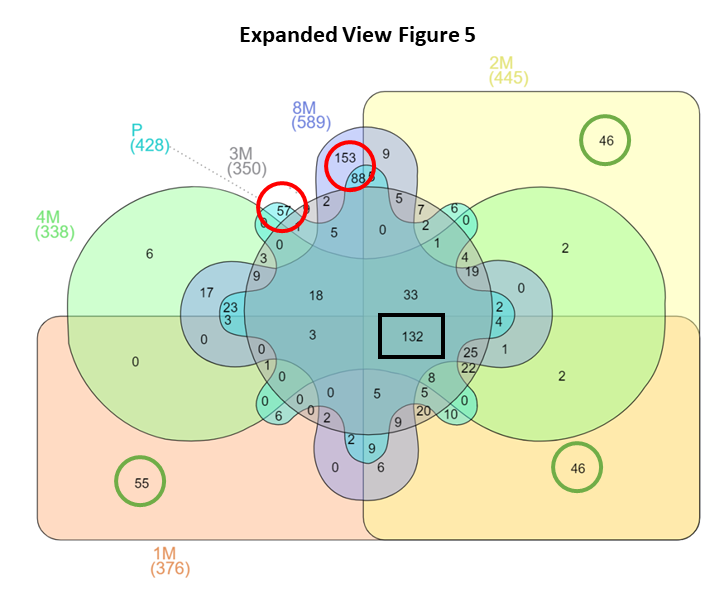

### Extended view figure 6

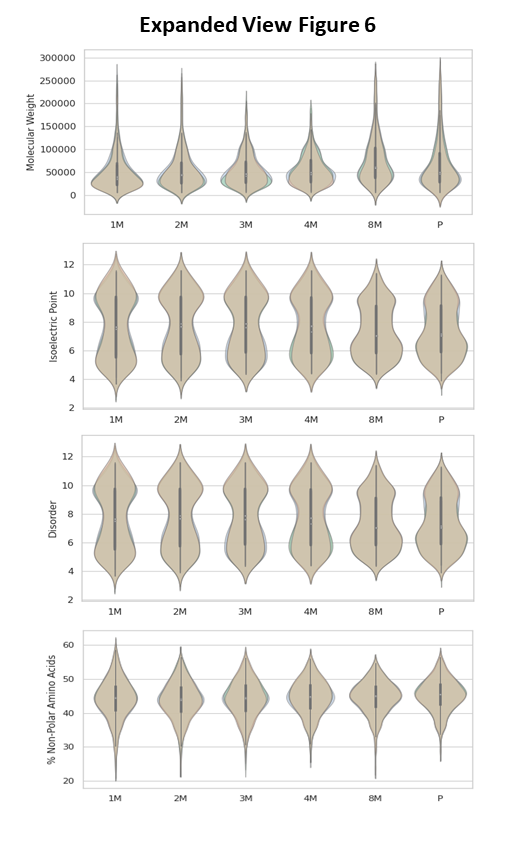

### Extended view figure 7

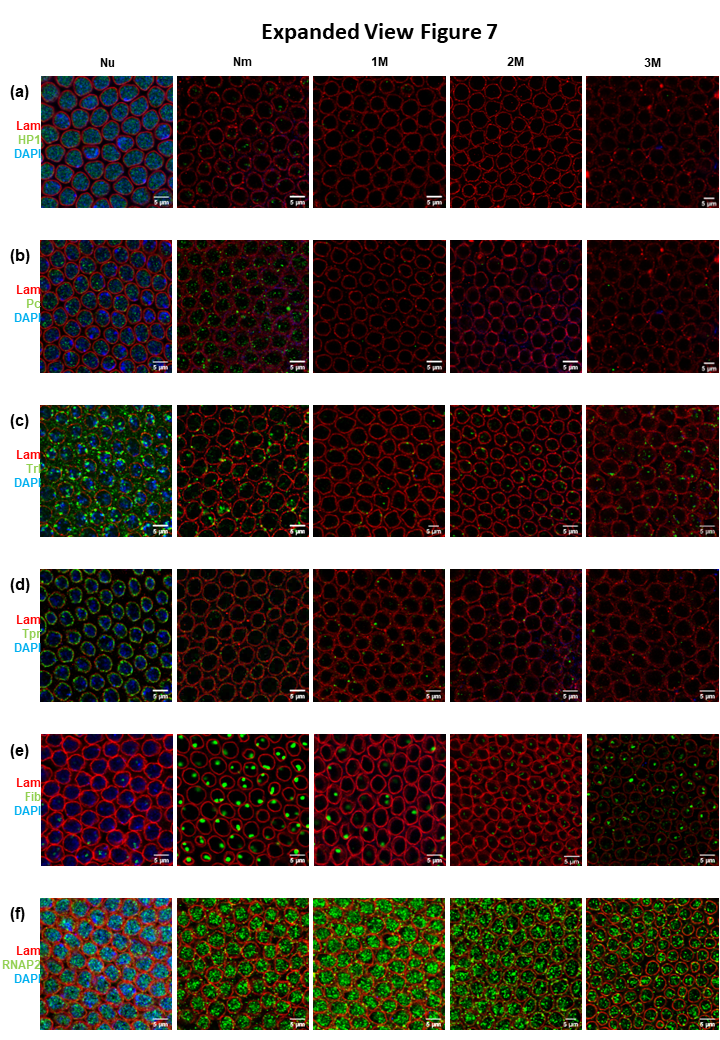

### Extended view figure 8

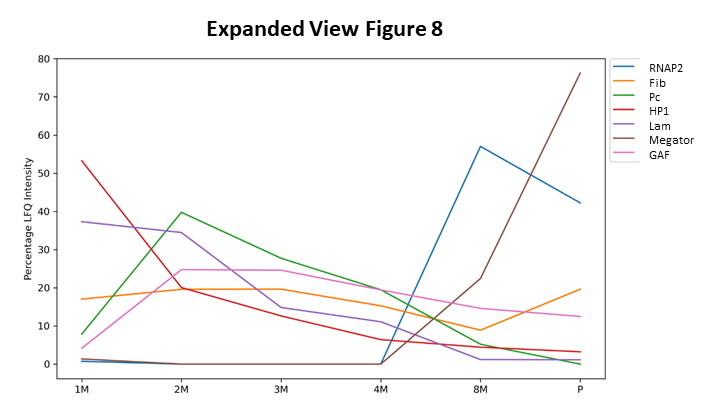

### Extended view figure 9

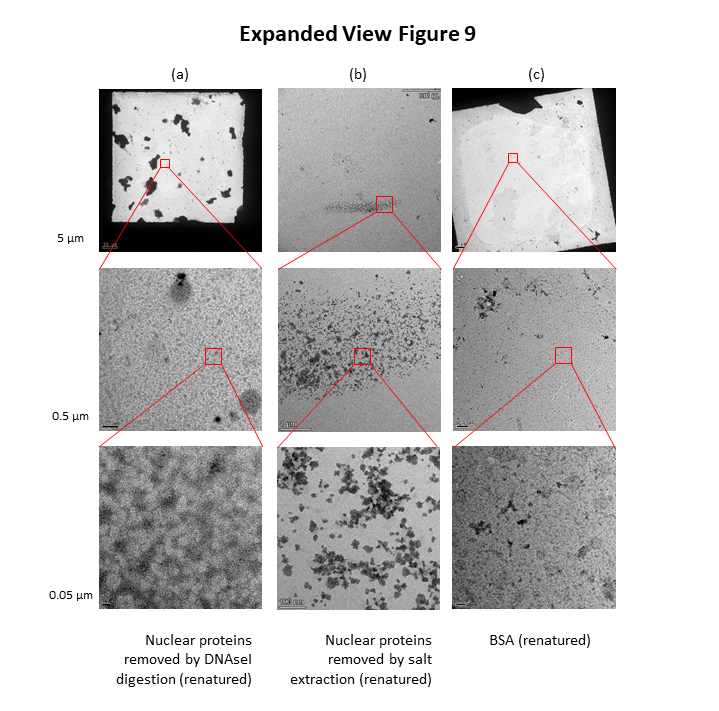

### Extended view figure 10

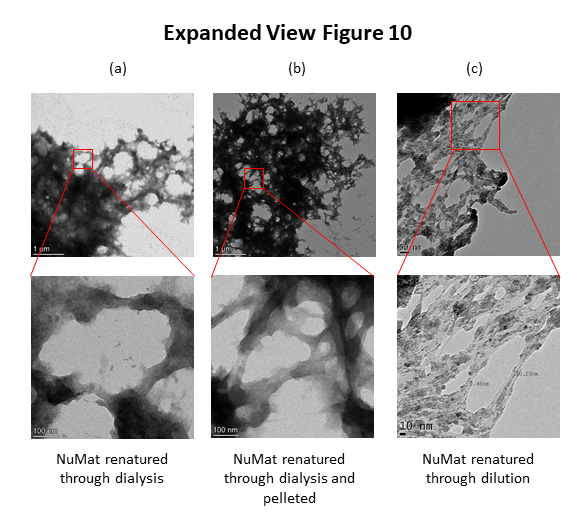

### Extended view figure 11

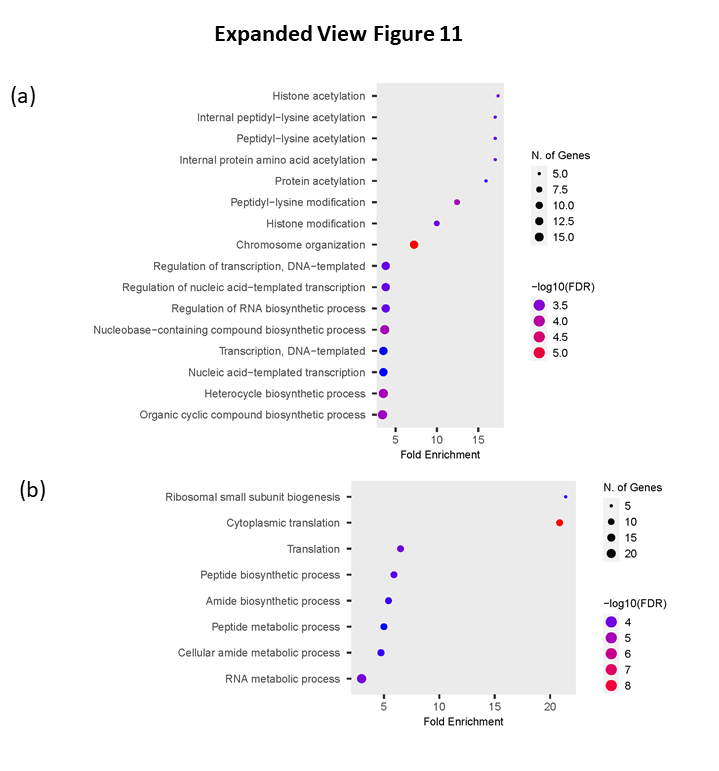
