## Supplementary material for "Biochemical deconstruction and reconstruction of Nuclear Matrix reveals the layers of nuclear organization": Supp file 1

**Supplementary material:**
The GdnHCl extraction process was standardized in several steps. To first assess which chaotrope suits the extraction of the NuMat, we performed parallel extractions with GdnHCl and Urea. Four separate NuMat aliquots were resuspended in 1 M, 2 M, 3 M, and 4 M GdnHCl solutions respectively, and yet another four were resuspended in 1.5 M, 3 M, 4.5 M, and 6 M Urea. These suspensions were incubated at room temperature for 20 minutes and then spun down. The supernatant and pellet were loaded on SDS-PAGE and visualized with silver stain. The profiles of the supernatant were markedly different from the profiles of the pellets, implying that the set of proteins solubilized in GdnHCl solutions was different from the ones retained. There is a gradual increase in band intensity from a lower concentration of GdnHCl to higher ones (Fig S2 a). On the other hand, no solubilization was seen with Urea (1.5 M, 3 M, 4.5 M, and 6 M) (Fig S2 b).

To standardize the ideal criteria for GdnHCl-based fractionation of NuMat, we first checked if proteins would keep leeching out in fresh buffers of the same concentration of GdnHCl. We sequentially extracted individual aliquots of NuMat with the same concentration of GdnHCl, switching the buffer every hour. For all four concentrations of GdnHCl considered, supernatant protein profiles show that most of the protein extractable at a concentration, is extracted within two hours (Fig S2 c). Having ascertained a lack of leeching, we sequentially extracted NuMat with increasing concentrations of GdnHCl (in order: 1M, 2M, 3M, 4M) by resuspending NuMat in 1M GdnHCl solution, incubating it for 2 hours, spinning it down, removing the supernatant and adding the extraction buffer with 2M GdnHCl, and so on for 3M and 4M GdnHCl extraction buffers. The profiles seem to be comparable in intensity – with different extraction patterns for different proteins (Fig S2 d). We observe three modes of extraction 1) extracted at low GdnHCl concentration or easily solubilized, 2) extracted at high GdnHCl concentration or difficult-to-solubilize, and 3) extracted equally at all concentrations.

GdnHCl has some known preferences for protein solubilization reported in the literature, showing a bias towards low molecular weight proteins of highly ionic and disordered nature (Möglich et al., 2005; Povarova et al., 2007, 2010). We plotted histograms for molecular weight, isoelectric point, intrinsic disorder fraction per protein, and percentage non-polar amino acids per protein (Fig S4). As shown, there seems to be no distinction of biophysical extractability in extraction by different GdnHCl concentrations.

Antibodies used for western blots

| **Antibody** | **Antigen from** | **Source** | **Accession number** | **Dilution** |
| --- | --- | --- | --- | --- |
| Lamin Dm0 | Fruit Fly | DSHB | ADL67.10 | 1000x |
| Pan H3 | Mouse | Abcam | Ab1791 | 5000x |
| GAPDH | Mouse | Abcam | Ab8245 | 2000x |
| Cytochrome C | Mouse | Abcam | Ab13575 | 1000x |
| Anti-Mouse HRP | Mouse | Abcam | Ab6820 | 10000x |
| Anti-Rabbit HRP | Rabbit | Abcam | Ab6802 | 10000x |
| Anti-Guinea-Pig HRP | Guinea Pig | Abcam | Ab6771 | 10000x |

Antibodies used for immunofluorescence

| **Antibody** | **Antigen from** | **Source** | **Accession number** | **Dilution** |
| --- | --- | --- | --- | --- |
| Lamin Dm0 | Fruit Fly | Dr. V. K. Parnaik lab | NA | 100x |
| BEAF | Fruit Fly | Raised in our lab | NA | 50x |
| HP1 | Fruit Fly | DSHB | C1A9 | 100x |
| Pc | Fruit Fly | SCBT | Sc-25762 | 50x |
| GAF | Fruit Fly | SCBT | Sc-98263 | 50x |
| Megator | Fruit Fly | DSHB | 12F10-5F11 | 100x |
| RNA pol2 CTD | Mouse | Abcam | Ab5131 | 100x |
| Fibrillarin | Mouse | Abcam | Ab5821 | 100x |
| Anti-Mouse DL488 | Mouse | Jackson | 715-485-150 | 500x |
| Anti-Rabbit DL549 | Rabbit | Jackson | 711-505-152 | 500x |
| Anti-Guinea-Pig AF647 | Guinea Pig | Jackson | 706-605-148 | 500x |

Crucial reagents recommended for reproducibility

| **Reagent** | **Company** | **Catalog ID** |
| --- | --- | --- |
| DNase I | Sigma | D4527 |
| GdnHCl | Sigma | G3272 |
| TEM Copper grids | EMS | CF400-CU |
| TEM Nickel grids | EMS | FCF200-NI |
